# A sensorimotor instability drives a locomotor transition during fish development

**DOI:** 10.1101/2025.10.10.681567

**Authors:** Monica Coraggioso, Leonardo Demarchi, Robert Wong, Vito Dichio, Chloé Chaumeton, Thomas Panier, Ghislaine Morvan-Dubois, Geoffrey Goodhill, Volker Bormuth, Georges Debrégeas

## Abstract

Animals rely on movement to survive — to explore their environment, find food and mates and avoid danger. During development, changes in body shape, muscle strength and physiological needs drive the continuous adjustment of locomotor patterns. How these changes are orchestrated in a flexible and adaptive manner remains unknown. We explore this question in *Danionella cerebrum*, a miniature freshwater fish that is emerging as an important vertebrate model in systems neuroscience. We identify a clear transition in locomotion, from continuous to burst-and-coast swimming occurring around 3 weeks of age. We demonstrate that this transition is an energy saving strategy, and that it reflects an instability in the sensorimotor process governing speed regulation. Rather than a preprogrammed developmental switch, it is therefore directly tied to the animal swimming strength. We confirmed this finding by manipulating sensory feedback in order to induce a similar transition at fixed developmental stages. Together, our results illustrate a dynamic interplay between body, brain, and environment during development, offering new insights into the principles governing adaptive locomotion.

## 1 Introduction

From birth to maturity, animal behavior adapts throughout development to accommodate changing body morphology and physiological demands. In many instances, the emergence of new behaviours is linked to brain development, driven by neuromodulatory signals or genetic programs. In mammals for instance, hormonal signalling and transcriptional dynamics shape cortical maturation, which in turn regulates behaviours such as reproduction [1]. In Drosophila larvae, neuropeptidergic signalling elicits new behaviours in preparation for metamorphosis [2]. In zebra finches, the brain appears to rely on genetically encoded predispositions to learn song tones, even when sensory input is absent or delayed [3].

Locomotion — the ability to move through the environment — is the behaviour most tightly linked to organismal design. It requires flexibility to accommodate morphological changes during growth [4], and adaptability to meet evolutionary pressures for efficient, rapid, and versatile movement [5]. At a fixed age, transitions between locomotor modes often reflect this dynamic interplay with the environment: in humans, the walk-to-run, or in horses, the trot-to-gallop switches have been interpreted as bifurcations between locomotor attractors, whose relative stability are shaped by both biomechanics and tasks demands [6–8]. Similar self-organizing principles emerge in robotics, where decentralized feedback alone can generate spontaneous gait transitions without preprogrammed rules [9]. How new locomotor modes emerge and start being executed during development to accompany body growth remains elusive. In particular, the relative contributions of neuronal maturation and biomechanics in driving these transitions are still unclear.

Addressing this question requires a model system that permits simultaneous monitoring of behavior and brain activity throughout development. *Danionella cerebrum*, a recently established vertebrate model, offers unique advantages in this regard [10, 11]. Its small, transparent brain enables longitudinal, whole-brain calcium imaging at cellular resolution [12–15], while its small size and stereotyped behaviors make it amenable to detailed behavioral analysis [16–18].

In this study, we combine high-speed behavioral monitoring, fluid dynamic simulations, mathematical modeling, and virtual reality experiments to investigate how *Danionella* locomotion adapts to body growth during its early life stages, from 2 to 5 weeks post fertlization (wpf). We reveal and quantitatively characterize a developmental transition occurring around 3 wpf, during which animals shift from continuous to intermittent swimming. We assess the consequence of this transition in terms of energetic efficiency, and then propose a mechanistic model to explain how intermittent swimming can spontaneously emerge. Finally, we discuss the functional advantage that this mechanism offers to trigger the transition at the appropriate developmental stage.

## 2 Results

### High-resolution tracking reveals developmental transition in swimming behavior

To understand how locomotion evolves during early development, we longitudinally tracked spontaneous swimming in *Danionella* from larval to juvenile stages using a custom high-speed tracking system (Fig. 1**a** and Methods). While the setup captures locomotion across multiple timescales (see Fig. 1**b**, SI Movie1 and SI Movie2), here we focus on fine-scale tail kinematics. We first quantified swimming by measuring each fish’s center-of-mass velocity, *v*_f_ (Fig. 1**c**). This simple metric is sufficient to reveal a clear developmental transition. Up to 2 wpf, swimming is predominantly continuous with only modest speed fluctuations. By 4–5 wpf, fish display intermittent locomotion, characterized by bursts of propulsion separated by inactive gliding phases. This shift emerges gradually, with both patterns coexisting around 3 wpf.

**Fig. 1.**
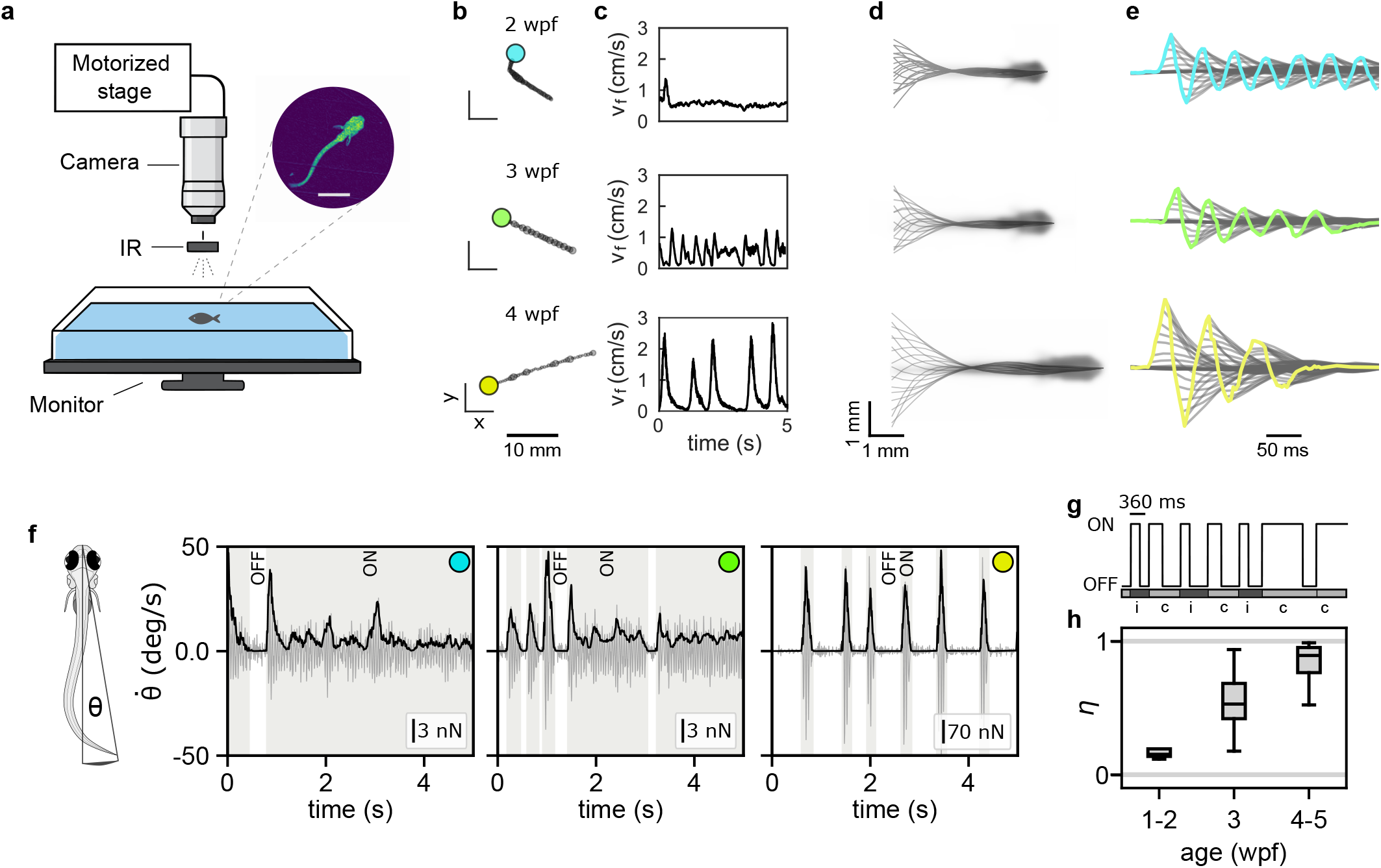
Behavioural tracking reveals continuous-to-intermittent transition during early development. **a**, Schematic of the experimental setup. A camera mounted on a motorized stage tracks the fish at 250 FPS. IR LEDs provide illumination from above while a monitor, positioned below the tank, displays a naturalistic background (stones). An example snapshot is shown on the top left. Scale bar: 1 mm. **b**, Five-second long trajectories for three example fish of 2 wpf (cyan), 3 wpf (green), and 4 wpf (yellow). **c**, Time evolution of centre-of-mass speed *v*_f_ for the same fish. **d**, Midline dynamics during forward motion. Time between consecutive profiles is 40 ms. The mean image of the fish across frames is shown in transparency. Scale bar: 1 mm. **e**, Midlines are time-shifted along the x-axis to visualize the time evolution of the tail tip (colored traces). **f**, Schematic definition of *θ*, the angle between the tail tip and the fish body axis, whose time derivative 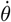 (in grey) is shown for three representative ages: 2 wpf (cyan), 3 wpf (green), and 4 wpf (yellow). The computed thrust signal (*F*_*T*_) is overlaid in black, and scaled to facilitate comparison between the two signal (the thrust scale is indicated by the inset bar). *F*_*T*_ is used to delineate ON and OFF states, represented respectively by the grey and white backgrounds. **g**, Swimming events are labeled as either intermittent (i) - if the corresponding ON state lasts less than 360 ms - or continuous swimming (c) - otherwise. Note that the threshold applies only to the duration of the ON state, but the event duration includes both the ON and subsequent OFF phases. **h**, Intermittency indices (*η*) for 3 age groups. The median and interquartile range (25th and 75th percentiles) are reported in the boxplot, with error bars indicating the full range of values.

We next asked whether this transition is reflected in the postural kinematics. Swimming involves axial muscle contractions that generate undulatory body waves increasing in amplitude from head to tail. Across ages (Fig. 1**d**), *Danionella* initiate lateral oscillations at a consistent trunk position, with a stable node located at 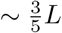 from the head tip, where *L* is the fish length. This suggests that the anterior-toposterior activation pattern is conserved. This finding is further supported by principal component analysis of segmented fish postures normalized by *L*. We found that the first three eigenmodes and their explained variance remaining consistent across ages (Fig. S1**a-d**). The characteristic tail-beat frequency also remains stable at *∼*20 Hz throughout development (Fig. S1**f**,**g**). Together these observations indicate that the postural kinematics at short timescales, *i*.*e*. on the order of the tail beating period, is conserved, with the motion of older and younger fish transformable into one another through simple length rescaling.

The developmental locomotor transition is thus primarly reflected in the slower temporal modulation of the tail-beat amplitude: tail-beating shifts from continuous oscillations to stereotypic discrete sequences of *≈* 3 cycles followed by periods of rest (Fig. 1**e** and Fig. S1**e**). To further characterize the emergence of burst-and-glide locomotion, we focused on the tail dynamics from which we extracted an effective thrust *F*_*T*_ using a fluid dynamic reactive model (see Methods). Compared to speed, *F*_*T*_ offers a better signal-to-noise ratio, enabling precise identification of motor activity periods: each time point was classified as either an ON state, marked by significant forward thrust, or an OFF state, corresponding to the absence of active movements (Fig. 1**f**). Early in development, the distributions of ON and OFF state durations are broad and approximately exponential, lacking a characteristic timescale. As the fish mature, they become narrower and develop a distinct peak, revealing stereotyped burst 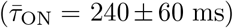 and pause 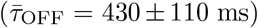 durations (see Fig. S2 and see Supplementary Information Section 1.1). To quantify this evolution, the thrust signals were segmented into swimming events (or bouts) labeled ‘intermittent’ if the ON duration was shorter than 360 ms—the mean plus two standard deviations of *τ*_ON_ in 4–5 wpf fish—and ‘continuous’ otherwise (Fig. 1**g**). We then computed an intermittency index *η* defined as the fraction of intermittent events among all swimming events.

With age, *η* increases from *η ≈* 0 (continuous swimming) to *η ≈* 1 (intermittent swimming), as shown in Fig. 1**h**. Notably, while early and late developmental stages show low variability in *η*, values at the intermediate stage (around 3 wpf) span the full range, reflecting the heterogeneous behavior during this transition phase (Fig. S3).

### The locomotor transition as an energy-saving strategy

We asked whether energy saving could be a key driver behind the developmental switch in *Danionella* locomotion. Continuous swimming is expected to be more efficient at small body sizes, whereas burst-and-glide can become advantageous as fish grow larger [19, 20]. As body length (*L*) and swimming speed (*v*_f_) increase, the inertial forces begin to dominate over viscous ones. In the inertial regime, interrupting tail motion reduces drag during glides, enhancing efficiency. The relative importance of inertial and viscous forces is captured by the Reynolds number, Re = *ρLv*_f_ */µ*, where *ρ* is the fluid density and *µ* its viscosity. In *Danionella*, We found that Re increases from 20 *±* 10 to 140 *±* 100 during the examined developmental window(Fig. 2**a-b**), with large intra-individual variability reflecting the broad distribution of swimming speeds, particularly in older fish.

**Fig. 2.**
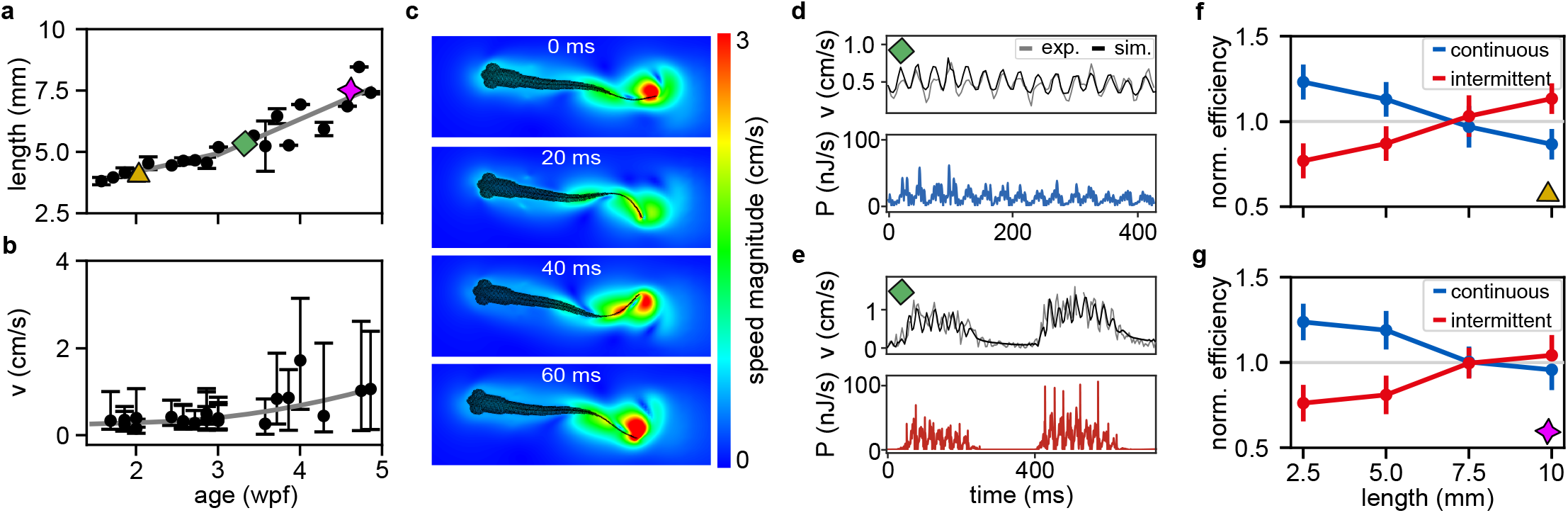
Hydrodynamics of continuous vs intermittent swimming. **a**, Fish length as a function of age (mean ± std across individuals). Symbols indicate the three 3D model fish used for CFD simulations. Lines show linear fits before and after 3 wpf: the growth rate increases from 0.1 (*≤*3 wpf) to 0.2 mm/day (*≥* 3 wpf). **b**, Median and interquartile range of fish speed during active swimming across development. **c**, Example 2D snapshots from CFD simulations, separated by 20 ms intervals. **d-e**, Simulated continuous (d) and intermittent (e) swimming events using the intermediate-age 3D model. Top: Simulated (black) and experimental (grey) center-of-mass velocities. Bottom: Dissipated power (*P*) predicted from simulations. **f**, Normalized efficiency vs. body length using the 14 dpf *Danionella* morphology. Mean *±* std across simulated events for continuous (blue, N = 11, 11, 11, 9) and intermittent (red, N = 14, 14, 14, 11) swimming. See Fig. S5**d** for non normalized data and statistical tests. **g**, Same analysis as in (f) using the 30 dpf morphology. Continuous (blue, N = 11, 11, 11, 9) and intermittent (red, N = 14, 14, 12, 13) swimming. See Fig. S5**e** for non normalized data and statistical tests.

To quantitatively assess how swimming energetics evolve with development, we performed computational fluid dynamics (CFD) simulations (see Methods) [21]. Realistic 3D *Danionella* models were reconstructed from anatomical scans (see Methods and Fig. S4) and animated using experimentally recorded tail kinematics (Fig. 2**c**, SI Movie3-6). To validate the CFD simulations, we computed the relative error between simulated and experimental speeds using an intermediate-age *Danionella*, performing both swimming modes (see Fig. 2**d–e** and Fig. S5). This data-driven approach enables direct comparison of energetic performance between continuous and intermittent swimming, consistent with a recent large-scale computational study that systematically evaluated the hydrodynamic efficiency of burst-and-glide gaits [22]. The simulated swimming motion generated fluid motion, which was used to compute the instantaneous dissipated power (*P*). Mechanical energy was obtained by integrating power over time, providing a proxy for metabolic energy cost. Efficiency was then estimated as the inverse of the energy per distance travelled.

To evaluate how swimming efficiency evolves during development, we ran CFD simulations using 3D *Danionella* models at 2 and 4 wpf, scaled to body lengths from 2.5 to 10 mm. At each size, we applied experimentally recorded tail kinematics to simulate both continuous and intermittent swimming. The resulting normalized efficiency is shown in Fig. 2**f–g**. The normalization factor, defined as the average efficiencies of continuous and intermittent swimming, enables a direct comparison between the two modes (see Fig. S5**d-e** for efficiency without normalization). In both morphologies, simulations revealed a size-dependent shift in energetic efficiency: continuous swimming is more efficient at smaller sizes, while burst-and-coast becomes favorable at larger sizes. The crossover occurs around 7 and 7.5 mm, slightly offset between the two 163 3D models, suggesting that body shape also modulates swimming efficiency. Notably, this size range (corresponding to *≈* 4 weeks-old fish) is slightly larger but close to the size at which the behavioral switch is observed, supporting the idea that burst-and-glide swimming emerges as a biomechanical adaptation to maintain energetic efficiency across development.

### The velocity overshoot at the onset of movement is predictive of the bout outcome

Having established the energetic advantage of transitioning from continuous to intermittent swimming, we now ask what mechanism triggers this transition in the developing fish. Since the selection between short and long swimming events must occur within the duration of a short bout, we focus on the speed signals immediately following movement onset. Figure 3**a** shows bout-averaged velocity traces aligned to movement onset. Fish were grouped into three categories based on their intermittency index *η*: continuous (*η <* 0.2), mixed (0.2 *< η <* 0.8), and intermittent (*η >* 0.8). In all groups, swimming begins with one to two high-amplitude tail oscillations that lead to a peak velocity (*v*_start_) reached after 100–150 ms. The fish speed then decays toward either a constant value or zero (see also Fig. 1**e**).

**Fig. 3.**
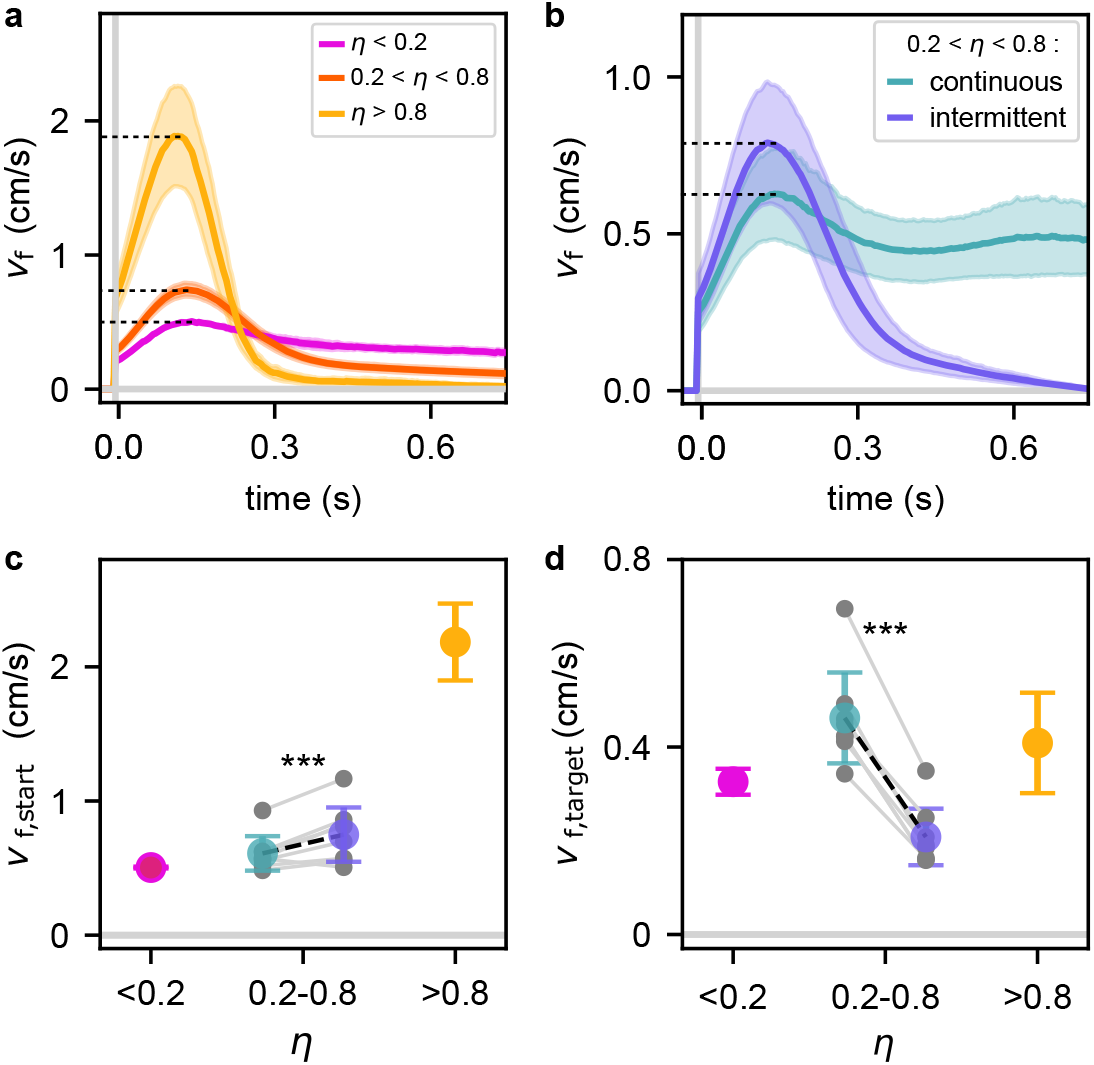
Velocity overshoot at movement onset differentiates swimming modes. **a**, Fish speed (*v*_f_) aligned to movement onset for forward swimming events, with the subsequent 750 ms of motion shown. For each individual fish between 10^2^ and 10^3^ such events were recorded, depending on whether their swimming was predominantly continuous or intermittent. Fish were grouped by intermittency index *η* into three categories: *η <* 0.2 (N = 4), 0.2 *< η <* 0.8 (N = 11), and *η >* 0.8 (N = 5). Bold lines show the mean across fish; shaded areas denote standard deviation. **b**, Same as in (a), restricted to the intermediate group (0.2 *< η <* 0.8). Events were color-coded as continuous (cyan) or intermittent (violet) based on the bout duration criterion defined in the main text. Bold lines and shaded regions represent the group mean and standard deviation, respectively. **c**, Overshoot maximum speed *v*_f,start_ as a function of *η*. Mean *±* std across fish are shown. In the intermediate regime (0.2 *< η <* 0.8), values are further separated into continuous (cyan) and intermittent (violet) events. Grey dots represent fish-specific means for each category. Two-sample t-test: *p* = 10^*−*4^. **d**, Same as in (c), for the target speed *v*_f,target_ (*p* = 10^*−*6^).

We found that *v*_start_ positively correlates with *η* (Spearman *r*_s_ = 0.71, *p* = 10^*−*4^, see Fig. S6). To further assess the predictive power of *v*_start_ on bout type outcome, we focused on the intermediate developmental stage (0.2 *< η <* 0.8), where both locomotor modes coexist. Swimming events were classified as either short (continuous) or prolonged (intermittent) using a duration threshold of *≈* 800 ms, defined as 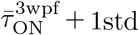 (Fig. 3**b**). The overshoot velocity *v*_start_ is lower in prolonged than in short bouts (Fig. 3**c**). This correlation was found to be statistically significant across tested fish in the intermediate regime (see Methods).

Overall, our analysis shows that the initial speed (reached within the first *∼* 100 *−* 150 ms after movement onset) largely determines the ensuing swimming pattern: high initial speed generally leads to short, rapidly interrupted bouts, whereas low initial speed tends to be followed by prolonged swimming events. As a consequence, although *v*_start_ grows by a factor of *≈* 4 across the continuous-to-intermittent transition, the average bout speed *v*_target_ increases by only *≈* 20% (see Fig. 3**c-d** and Fig. S6b-c).

### Instability induced by sensorimotor delays drives swimming mode transition

In this section, we build on the previous observations to propose a mechanistic model that could account for the transition to intermittent swimming. Our hypothesis is that the emergence of this new locomotor mode is due to a dynamic instability of the sensorimotor system used by the animal to regulate its speed.

During continuous swimming, we observed quasi-periodic speed oscillations with a period of *≈* 400 ms (see Fig. S8), reminiscent of those reported when *Danionella* larvae regulate their swimming speed in a virtual environment [15]. In that context, we showed that these oscillations arise from a finite delay (*≈* 100 ms) in the visuomotor feedback loop. This suggests that *Danionella* rely on sensory feedback for speed regulation even during freely swimming, and that similar delays influence this control. Such a mechanism can also explain the velocity overshoot at bout onset (Fig. 3**a–b**): the initial high velocity period likely reflects a ballistic phase (no feedback) lasting for *≈*100 ms, followed by deceleration as sensorimotor regulation engages to drive the system toward a target speed.

The sensorimotor loop is schematized in Fig. 4**a**. We assume that the fish can estimate its speed *v*_f_ (the way it does so will be discussed in the next section) and adjusts its motor command *u* based on the difference between *v*_f_ and an internally defined target speed *v*_f,target_. This motor command drives the tail oscillation speed *v*_t_, which in turn sets *v*_f_. The loop can be mathematically described by a system of three linear equations:

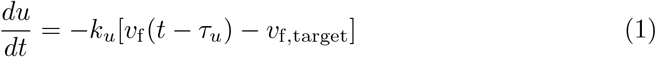

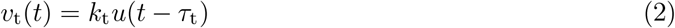

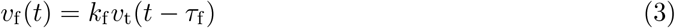

where *v*_t_ is the tail-tip speed averaged over one oscillation cycle: *v*_t_ = 2*Af*, with *A* the peak-to-peak amplitude and *f* the (invariant) tail-beat frequency. The motor command *u* denotes an upstream neuronal signal driving the tail amplitude. Each step in the sensorimotor loop has an associated gain (*k*_*u*_, *k*_*t*_, *k*_*f*_) and delay (*τ*_*u*_, *τ*_*t*_, *τ*_*f*_). Although sensory and motor signals are likely encoded nonlinearly in sensorimotor circuits [15], here we assume linear relationships in Eq.(1) and Eq.(2) for interpretability; however, we show in Supplementary Information Section 1.3 that including logarithmic nonlinearities yield analogous conclusions.

**Fig. 4.**
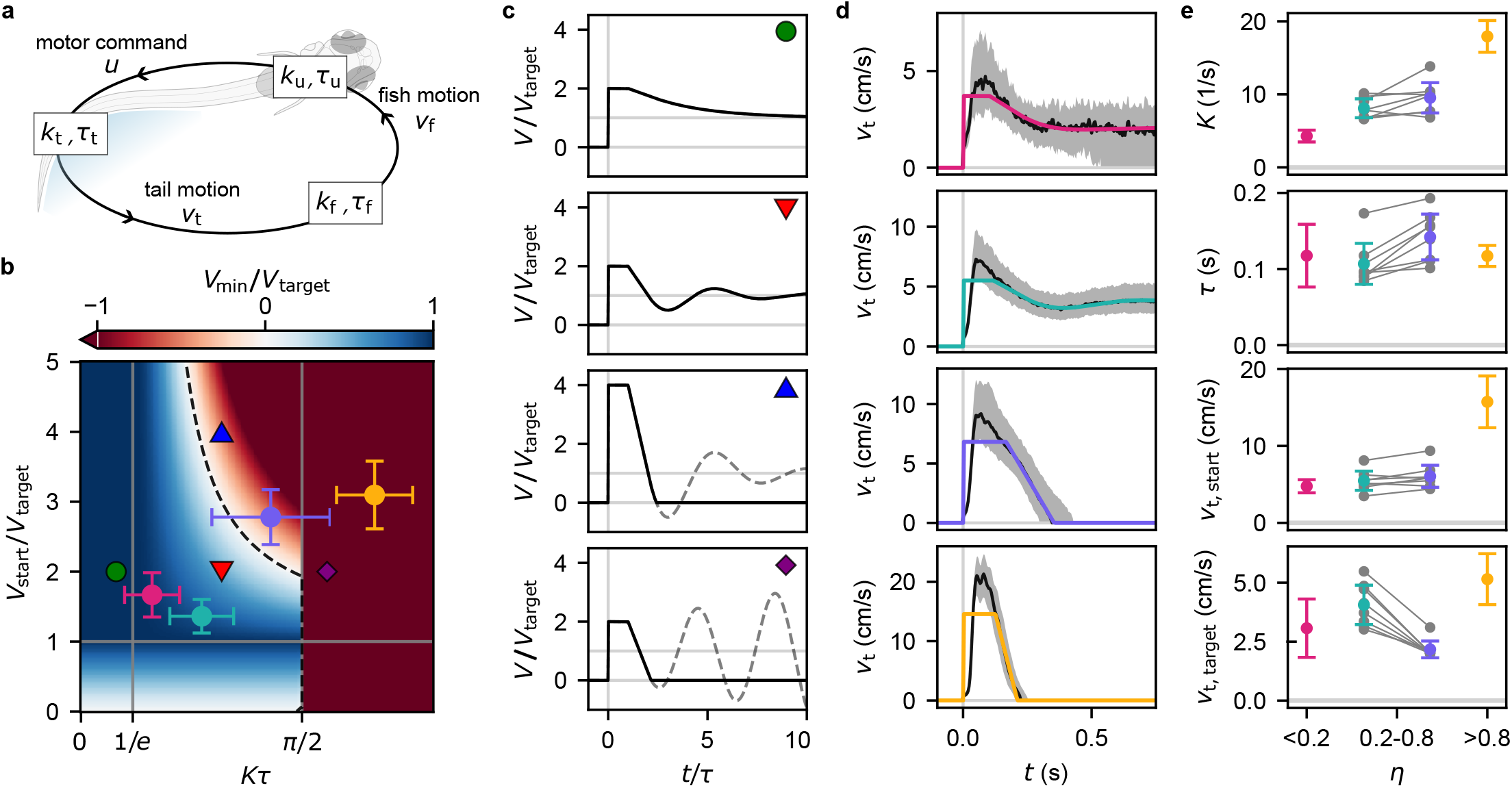
A model with delayed feedback control reproduces the locomotor transition. **a**, Schematic representation of the sensorimotor loop for speed regulation. **b**, Minimum speed *V*_min_ computed from Eq. (S3) as a function of the gain delay product *Kτ* and the ratio of initial to target speed *V*start*/V*target. The grey solid lines delineates the different dynamic regimes in the phase space. The black dashed line corresponds to *V*_min_ = 0 and separates the regions of continuous (blue) and intermittent (red) swimming. **c**, Example traces showing the evolution of the speed according to Eq. (S2) for different values of the parameters (corresponding symbols in b). If the speed reaches zero, swimming stops (black solid line), even though the mathematical solution would continue to oscillate (grey dashed line). **d**, Fitting Eq. (S2) (colored solid lines) on the speed onsets *v*t(*t*) (black solid lines and grey shaded areas indicate medians and interquantile ranges) for three example fish used in the computation of Fig. 3**a-b. e**, Best fit values of the parameters shown for different fish (errobars indicate mean and standard deviation), separated in four groups as in Fig. 3**c**. These values were used to compute the corresponding errorbars in panel b. The values corresponding to individual fish are shown as grey points for 3 wpf fish.

Eq.(3) describes how tail motion determines the fish’s forward swimming speed. The proportionality constant *k*_f_ can be expressed in terms of the Strouhal number St = *Af/v*_f_ as *k*_f_ = 1*/*(2St). Consistent with literature [23], we found that St slowly decreases with Re, resulting in a small increase of *k*_f_ across the probed developmental window (see Fig. S9). Regarding the delay *τ*_f_, we found it to be approximatly invariant. It should be noted however that the sharp delay used in Eq.(3) is a simplifying approximation as an exponential kernel provides a better match between tail and fish speed.

Equations (1-3) can be combined into a single one for *u, v*_t_, or *v*_f_, which we write generically for a variable *V* as:

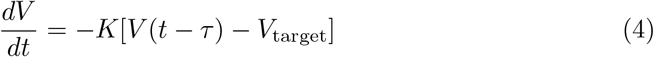

where we have introduced the total delay *τ* = *τ*_u_ + *τ*_t_ + *τ*_f_ and gain *K* = *k*_u_*k*_t_*k*_f_ in the sensorimotor loop. Eq.(4) is a linear delayed feedback equation which can be solved analytically [15]. Its behavior depends on the sole dimensionless parameter *Kτ* (see Supplementary Information Section 1.2). For *Kτ <* 1*/e*, the solution converges exponentially to *V*_target_, for 1*/e < Kτ < π/*2, it converges to the same value through damped oscillations, while for *Kτ > π/*2, the system is unstable and displays diverging oscillations (Fig. 4**b-c**). Assume a fish starts swimming with an initial speed *V*_start_. After a time *τ*, it begins regulating its speed toward the target value *V*_target_. Depending on the gain-delay product *Kτ* and the ratio *V*_start_*/V*_target_, the speed may either converge to the target value — possibly with damped oscillations — or it can reach a negative minimum value *V*_min_ (Fig. 4**c**). Since negative speeds are physiologically impossible, the fish effectively halts, resulting in the execution of a short bout. This bifurcation, inherent to the delay differential equation that governs sensorimotor regulation (Eq. (4)), thus offers a simple mechanism for the observed locomotor transition.

We sought to infer the model parameters values for fish at various developmental stages. To this end, we fitted the evolution of the tail speed at the onset of swimming using Eq.(S2) (see Methods). For fish in the intermediate stage (0.2 *< η <* 0.8) we separately considered the short and prolonged bouts as in Fig. 3**b**. The model reproduces the observed velocity decay across conditions, preceded by an initial period *τ* during which we assume constant *v*_t_ (Fig. 4**d**). The initial and target tail speeds *v*_t,start*/*target_ estimated from the fit (Fig. 4**e**) follow the same trends as those obtained from the measured fish speed in Fig. 3**c–d**. The infered sensorimotor delay *τ* remains stable across fish and ages, while the gain *K* increases *≈* 4-fold with development. When represented in the dynamic phase space, we find that, as fish grow, the system moves into the unstable region (Fig. 4**b**). Variability in swimming mode at the intermediate stage appears to be largely explained by differences in initial and target speeds: higher *v*_t,start_*/v*_t,target_ ratios lead to short bouts, lower ones to continuous swimming.

As *τ* is approximately constant and *v*_t,target_ varies weakly across development, the transition to instability can be primarily attributed to the growth in *K* and *v*_t,start_. We recall that the sensorimotor gain *K* = *k*_*u*_*k*_t_*k*_f_ encompasses the three successive transformations of the sensorimotor loop. Since *k*_f_ shows only a moderate developmental increase, this cannot explain the observed fourfold rise in *K*. Furthermore, within the scope of this model, the transition is driven by morphological rather than neuronal cues. Thus, we assume that the responsiveness *k*_u_ (and the initial motor command *u*_start_) remains unchanged across development. Under these hypotheses, the transition is mainly driven by *k*_t_. This parameter, which sets the relationship between motor command and tail speed, is indeed expected to increase as the fish grows and progressively develops more strength and greater thrust [24]. This increase in *k*_t_ concurrently drives an increase in the initial speed *v*_t,start_ = *k*_t_*u*_start_ for a given motor command.

We conclude that, while maintaining the motor control mechanism for speed regulation unchanged, the sole strengthening of the fish during development can drive the system across the bifurcation threshold, corresponding to a transition from continuous to intermittent swimming.

### Transition between continuous and intermittent swimming can be externally induced

Our mechanistic model implies that *Danionella* can internally estimate self-motion — with a short delay — in order to gauge its own speed *v*_f_ and achieve a target speed *v*_f,target_. Self-motion estimation is a complex and distributed multisensory process that remains challenging to dissect. Here we aim to identify which sensory modality dominates in freely swimming fish and then test the model by manipulating sensory feedback to assess its impact on the swimming mode selection.

We ran experiments in which we successively altered the visual, lateral line and vestibular feedback in freely swimming animals. We found that locomotor behavior was not significantly modified in fish swimming in complete darkness or after lateral line ablation (see Methods and Fig. S10), indicating that neither vision nor the lateral line system serves as the primary input for self-motion estimation. To externally manipulate vestibular input, we placed 5 wpf *Danionella* in dextran solution with a viscosity ten times higher than that of water (see Methods). We observed that the fish, which at this age swim intermittently in water, now reversed to continuous swimming as shown by the large increase in the swim event duration (Fig. 5**b** and SI Movie7). This observation is consistent with the model prediction under the assumption that velocity estimation relies primarily on vestibular input. Indeed, the increase in medium viscosity leads to an effective reduction of the coefficient *k*_f_ that controls the relationship betweeen the fish speed and the tail motion, as evidenced by the *≈* 3 fold reduction of the onset speed *v*_f,start_ (Fig. 5**a**). As a consequence, the system is expected to cross back over the bifurcation line and settle in the continuous regime of the phase space, corresponding to low *Kτ* and low *v*_f,start_*/v*_f,target_ (see Fig. 4**b**).

**Fig. 5.**
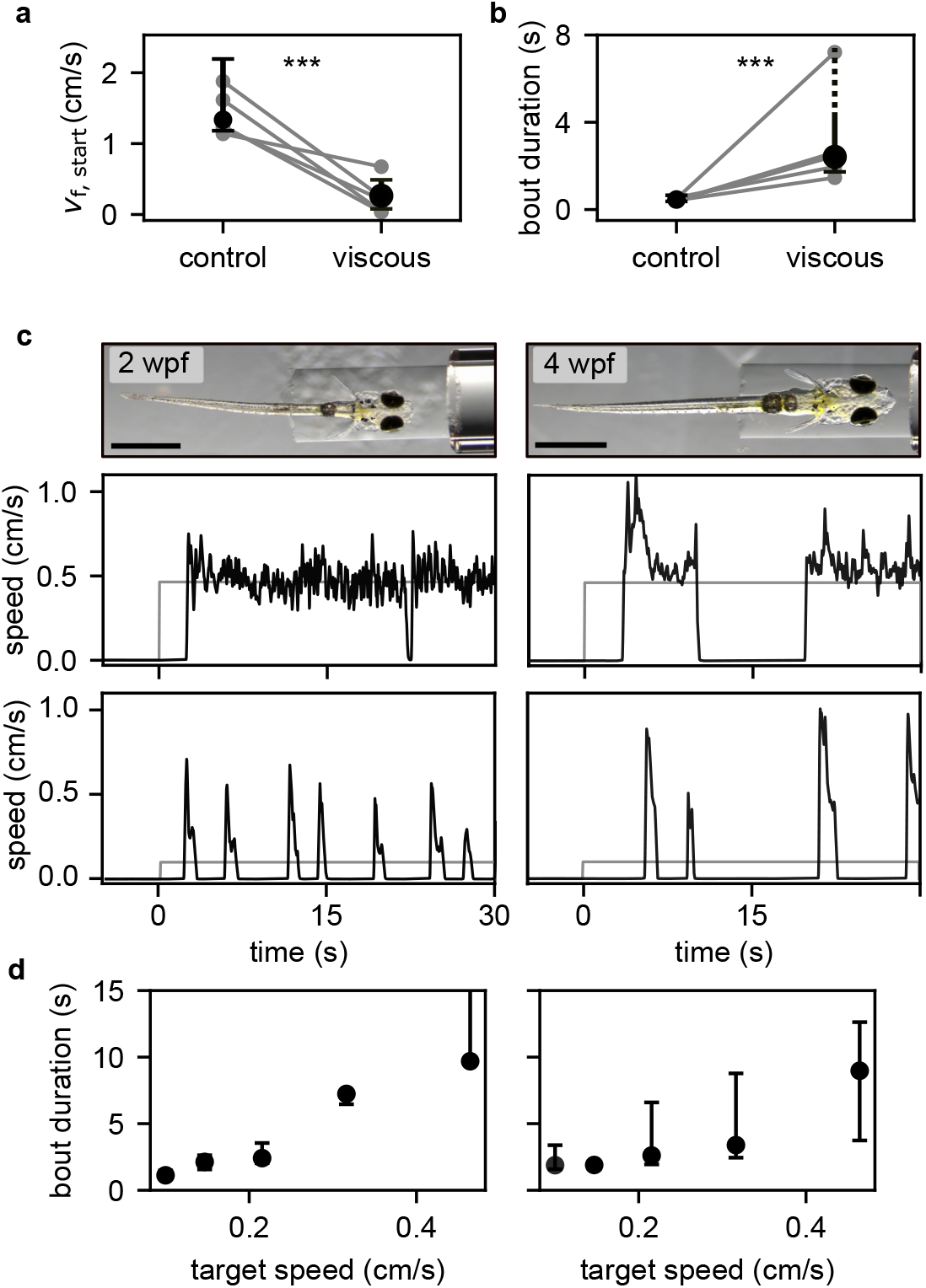
Locomotor transition can be externally induced. Comparison of swimming kinematics in water (control) and viscous medium. **a**, *v*_f,start_ (median ± IQR) across fish (N = 5); grey lines show average values of different swimming events for individual fish. Mann–Whitney U test: p = 10^*−*4^. **b**, Bout duration (median ± IQR) across fish. In viscous media, fish typically swam until reaching the end of the - smaller - experimental tank (see Methods), making the upper bound of bout duration not meaningful. Mann–Whitney U test: p = 10^*−*4^. **c**, Top: head-fixed fish at 2 and 4 wpf, prepared for virtual reality experiments. Scale bar: 1 mm. Bottom: example traces of fish fictive velocity (black) in response to forward visual motion for two target speeds (shown in grey). **d**, Bout duration (median ± IQR) versus target speed in 2 wpf (left, N = 5) and 4 wpf (right, N = 5) fish (N = 5 each).

We wondered whether this effect is a specific byproduct of altered vestibular input, or whether it more generally reflects the system response to changes in the sensorimotor loop. To this end, we performed experiments in head-tethered animals (Fig. 5**c**), in which vestibular feedback is absent, using the same virtual reality setup as in Demarchi *et al*. [15] (see Methods). In this configuration, the fish relies on optic flow to regulate its speed and stabilize position in response to externally imposed visual currents, a behavior known as the optomotor response. While in the freely swimming configuration, the target speed is set internally, here we can externally define *v*_t,target_ and probe its effect on the swimming regime (see also Supplementary Information Section 1.3). In 2 wpf fish, lowering the target speed triggered a shift from continuous to intermittent swimming (Fig. 5**d**, left). Conversely, we found that 4 wpf fish could switch to continuous swimming for high enough target speed (Fig. 5**d**, right). Thus, for both ages, bout duration increased with target speed (Fig. 5**e**; see also Fig. S3**b** for bout durations in freely swimming fish).

Altogether, these results show that *Danionella* can engage in either swimming modality at any developmental stage, and that the selected swimming pattern depends on the sensorimotor system and can be externally controlled.

## 3 Discussion

By systematically monitoring the maturation of locomotor behaviour during early development in *Danionella cerebrum*, we identified a transition from continuous to burst-and-glide swimming that occurs at around 3 weeks of age. We observed that the acceleration at the onset of movement is predictive of the ensuing swimming modality: high initial acceleration tends to be followed by a sharp deceleration and the interruption of movement, whereas low acceleration is likely to initiate a prolonged swimming period. This non-trivial correlation supports a mechanistic model of sensory-regulated swimming in which the locomotor transition naturally emerges as a dynamic instability of the delay sensorimotor loop through which the animal regulates its speed.

The model accounts for the observation that fish in the intermediate regime can exhibit both swimming modes, as they can initiate movements with acceleration above or below the threshold of instability. It also explains why intermittently swimming fish placed in a viscous fluid revert to continuous navigation. Indeed, within the scope of this model, the locomotor modality is determined by the hydrodynamic regime rather than by chronological age.

The sensorimotor feedback loop underlying the locomotor instability requires selfmotion estimation. To compute this estimate, animals may use visual, proprioceptive, mechanosensory, motor efference copy, and vestibular inputs. Our experiments show that freely swimming *Danionella* primarily estimate their own speed by integrating vestibular signals at movement onset. However, on a longer timescale, this estimated velocity will accumulate errors, so we expect other sensory modalities to contribute to speed regulation, as observed in mice [25] and Drosophila [26]. This is also the case when vestibular feedback is unavailable, as in head-tethered configurations, for which *Danionella* rely on optic flow to regulate their speed.

Our hydrodynamic simulations showed that the continuous-to-intermittent transition enables the fish to minimize energy expenditure across development. Consistent with previous studies [19, 20, 22, 27–29], burst-and-glide swimming becomes energetically favourable as inertial forces become more important during growth. Based on energetics alone, one would however expect the transition to occur at 4 rather than 3 weeks of age. This small discrepancy in the transition timing could reflect the fact that energetics alone cannot fully explain the complex ecological and evolutionary adaptations of aquatic organisms across body sizes. For instance, oxygenation or self-generated sensory noise may depend on the swimming mode and could also play a role in determining the optimal swimming strategy [30, 31].

The proposed mechanism offers a simple, adaptive, and flexible control strategy to trigger locomotor transition. In our model, the internal sensorimotor parameters (notably *k*_*u*_ and *u*_start_) are evolutionarily tuned so that the transition is triggered when intermittent swimming becomes energetically beneficial. Once these parameters are set, the switch becomes solely dependent on the fish physiology through the thrust that the animal can develop, and not tied to a rigid developmental program. This mechanism ensures that the transition timing is robust against the large variability in body growth observed in natural conditions due to, e.g., differences in rearing temperatures [32] or sexual specialization [33].

In this simplified modeling framework, the motor output signal is reduced to a single parameter that controls the amplitude of tail oscillations. This parameter should therefore correspond to the activity of an upstream circuit regulating tail-beat amplitude. In zebrafish, the medial longitudinal fasciculus (nMLF) has been identified as a key higher-order center to control the animal forward speed [34, 35]. In *Danionella* larvae, this midbrain neuronal population has been shown to similarly correlate with the animal swimming speed [14, 15, 30], making it a natural candidate to encode the output signal of the sensorimotor process.

Given the morphological and ecological proximity between larval zebrafish and *Danionella*, one may ask whether the same mechanism could be at play in zebrafish larvae. From the onset of motility, zebrafish swim intermittently, and unlike *Danionella*, we were unable to induce a switch to continuous swimming by altering medium viscosity (data not shown). Similarly, during OMR assays, zebrafish consistently swam intermittently, regardless of the visual target speed [34, 36, 37]. In *Danionella*, continuous swimming occurs over a relatively short developmental window (1-3 weeks) during which the animal length increases by less than 20% (Fig. 2**a**, growth rate ≈ 0.1 mm/day). In contrast, zebrafish grow steadily up to the juvenile stage (growth rate *≈ 0*.*1 ≈* 0.2 mm/day [38]), and based on energetic considerations alone, one would expect an even shorter period of continuous swimming. Other constraints, beyond energetics, may have evolutionary driven the complete elimination of this locomotor pattern. The impossibility to induce continuous swimming through sensory manipulation in zebrafish actually suggests that intermittent motion is hardwired in this animal. These behavioral differences point to potential species-specific solutions to the form-function optimization problem, opening avenues for cross-species comparative studies.

Extending to artificial systems, the sensorimotor adaptive mechanism we propose could inspire the design of self-adjusting robots capable of switching between swimming modes based on global parameters to optimize efficiency [39].

## 4 Methods

### 4.1 Fish care

All experimental procedures involving fish (*Danionella cerebrum*) were approved by the ethics committee “Le Comité d’Éthique pour l’Expérimentation Animale Charles Darwin C2EA-005” (APAFIS authorization number: 52420-2024121210531432 v7). Fish are housed in the facility, with water temperature maintained at approximately 27 ^*°*^C, pH at *∼* 7.3, and conductivity around 330 µS cm^*−*1^. The room lighting followed a 14/10-hour day/night cycle.

Tanks were monitored daily to collect newly laid egg clutches, which were transferred to Petri dishes containing E3 medium and kept at 28 ^*°*^C in an incubator. After hatching, larvae were transferred to 1 L beakers and fed with rotifers twice a day until two weeks of age (±1 day), after which they were either used for experiments or transferred to 3 L tanks in the main water flow system. Adult fish were maintained in a recirculating water system, housed in groups of approximately 50 individuals in tanks with a volume of 30 L.

All the experiments were conducted using wild-type animals. Animals were transferred 14 from the fish facility to the experimental room one day prior to the planned experiments. All experiments were performed between 10:00 am and 5:00 pm. Fish were euthanized immediately following the conclusion of the experiments.

### 4.2 Swimming assay design

Individual fish swam freely in a large 40 *×* 40 cm^2^ arena (approximately 100*×* the fish length), filled with filtered system water to a depth of 1 cm. This design (see Fig.1**a**) minimizes wall effects and promotes natural exploratory behavior.

A camera (FlyCapture), mounted on a motorized stage, followed the fish in real time and recorded images at 250 FPS. A 3D printer machine (Carbide 3D Shapeoko 4) was repurposed to house both the camera and infrared illumination (LED and beamsplitter Thorlabs). Online tracking was optimized for accuracy using the YOLO computer vision model [40], implemented via the Ultralytics Python library, and for speed through parallel computation using Python’s built-in threading module.

Each experiment had a nominal duration of 2 hours. An agar frame was placed around the arena to prevent tracking failures near the edges. If tracking failed, the fish was re-centered and the experiment restarted with a new run. For data analysis, multiple runs from the same fish were pooled together. Experiments were conducted under controlled conditions at a water temperature of 26 ^*°*^C and constant illumination.

To minimize disturbances, visual stimuli were reduced using a dark infrared filter (Goodfellow) shielding the motorized stage, and vibrations were limited by mechanically decoupling the stage from the arena.

The overall setup was inspired by Johnson *et al*. [41], with improvements to enhance spatiotemporal camera resolution and tracking efficiency. Hardware and software were customized for these experiments, and programming was implemented in Python.

### 4.3 Pose estimation and kinematic extraction

The pose of the animal was extracted from raw video recordings using a custom Python-based workflow. Initially, we applied SLEAP pose tracking [42] to detect 5 keypoints along fish body, while also identifying and discarding low-confidence frames based on the model’s confidence scores. To further refine positional accuracy, each frame was registered using a reference image of the head to correct for residual rigid body transformations. Finally, the midline of the fish was segmented in 20 equispaced points using the Stytra algorithm [43]. Since postures are measured in 2D projection, the body length varies slightly across frames due to changes in animal pitch angle; therefore, in Fig. 2**a**, *L* is reported as mean *±* standard deviation rather than a fixed value.

All video-derived measurements are transformed into the laboratory frame of reference by appropriately combining them with the motor position recorded during the experiment. The instantaneous speed of the fish is given by 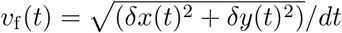, prior Savgol filtering of *x*(*t*) and *y*(*t*) in a time window of 60 ms to filter out ondulations of the body center of mass.

The speed of the tail is defined as 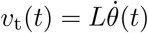, where *L* is the fish length and *θ* is the angle formed between the tip of the tail and the fish midline. In particular, we compute 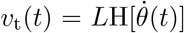, where 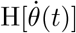 is the Hilbert transform of 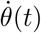, used to extract its envelop.

### 4.4 Reactive model for thrust estimation

To estimate the thrust from the tail movements we followed an approach that was used in previous studies on undulatory locomotion at intermediate Reynolds numbers [44]. We define a curvilinear coordinate system along the tail, with *s* going from 0 at the tip of the head to *l* at the tip of the tail *l*. If 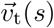 is the speed of the tail element at position *s* and 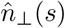 the unit vector perpendicular to the local tail segment, then 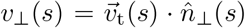 is the component of the speed perpendicular to the tail itself. Then, the reactive force on a tail segment of length *ds* at position *s* along the tail is given by:

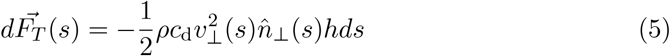

Where *c*_d_ is the drag coefficient, *ρ* is the water density, and *h* is the height of the tail segment. Then, if the fish heading direction is along 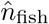, the instantaneous thrust resulting from the motion of the segment is given by the component along the direction of motion 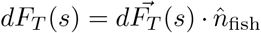. We get the total average thrust by integrating over the whole tail and averaging over one tail-beat period:

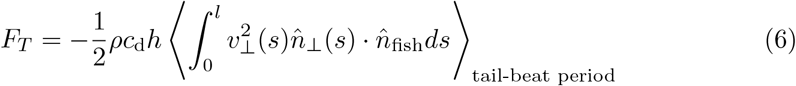

To compute *F*_*T*_, we considered an approximately constant tail height *h ≈* 0.7 mm and drag coefficient *c*_d_ *≈* 1.

Using *F*_*T*_, we defined ON and OFF states by setting a threshold that maximally separates active tail movements from background noise.

### 4.5 Computational fluid dynamics simulations

The CFD simulations were adapted from previous immersed boundary frameworks [21, 45]. At each time-step, experimentally recorded midline kinematics were mapped onto a 3D fish surface model to generate time-resolved surface velocities. Relative x–y displacements of each midline point were used to drive the motion of corresponding transverse slices of the 3D mesh. Velocity mapping and preprocessing were implemented using the Eigen C++ library [46].

Simulations were run in IBAMR (v0.15.0 and v0.16.0) with native MPI support for parallel computation. A domain-refined adaptive grid was used, and simulation parameters (e.g., viscosity, density) matched experimental estimates. Validation followed previous approaches [21], comparing CFD-predicted speed at the swim bladder with experimental measurements.

Energy was calculated as mechanical power output integrated over time, and efficiency was defined as distance traveled per unit energy. To compare swimming styles, we used tail kinematics recorded from 13 and 34 dpf *Danionella* as representative examples of the two extreme phenotypes: continuous and intermittent swimming, respectively. These kinematics were applied to animate 3D model fish at 14 and 30 dpf, rescaled to four different body lengths (2.5, 5, 7.5, and 10 mm). The rationale was not only to test the effect of body length on swimming energetics in the two swimming modes, but also to assess the contribution of morphological changes in body shape between 2 and 5 weeks.

Simulation results were visualized and post-processed using VisIt [47].

#### 4.5.1 3D fish models

3D fish models were obtained from lightsheet-based 3D stacks of clarified specimen, using the CUBIC-R+(N) clearing protocol from [48]. After fixation and fluorescein fluorescence staining, samples were embedded in low gelling temperature agarose (2%; A9414, Sigma-Aldrich). They were then incubated in 2 mL of 50% CUBIC-R+(N) reagent, composed of 45% (w/w) antipyrine (A5882, Sigma-Aldrich), 30% (w/w) nicotinamide (A15970, Thermo Fisher), and 0.5% (w/w) N-butyldiethanolamine (471240, Sigma-Aldrich), diluted in MilliQ water. Incubation was performed for one day at room temperature (RT), followed by one day in 2 mL of 100% CUBIC-R+(N). Specimen were then transfered in fresh CUBIC-R+(N) reagent inside the imaging tank, and left for a few minutes to allow refractive index homogenization.

**Table 1.** Optical configuration of the lightsheet system.

| System | Type | Objective (NA) | Lasers |
| --- | --- | --- | --- |
| Lightsheet Alpha3<br>PhaseView, France | Lightsheet | XLPLN 10X Olympus objective (0.6 NA), multi-immersion refractive index adaptive collar (RI 1.33–1.52) | 405 - **488** - 561 - 635 nm |

Lightsheet imaging was performed with the Alpha3 system (see tab.1), equipped with a 10× XL Plan N objective (XLPLN10XSVMP, Olympus, Japan) with a refractive index adaptive collar (RI 1.33–1.52), and an sCMOS Orca Flash4 camera (Hamamatsu, Japan). The acquisition software QtSPIM (version 2.20) was used. A 488 nm excitation laser and the ET525/50m emission filter were used for imaging. Lightsheet images were converted using arivis SIS Converter 3.1.1 and analyzed with arivis Vision4D 3.0.1 software (arivis AG, Germany).

### 4.6 Correlation between onset initial speed and bout duration

In order to quantify th ecorrelation between the onset initial speed *v*_*start*_ and the swimming event duration, we computed receiver operating characteristic (ROC) curves for all intermediate-stage fish (Fig. S7). For all but two animals, the ROC deviated significantly from chance (z-score *>* 3*σ*), with a mean area under the curve AUC_ROC_ = 0.63 *±* 0.08, indicating moderate but significant predictive accuracy.

### 4.7 Fitting onsets with delay equation

For each fish, we considered the onset speed signal averaged over all swimming events. We focused on the tail speed signal *v*_t_ as we expect it to more closely reflect the dynamics of the underlying motor command. We fitted the signal using Eq. (S2) (i.e. the solution of Eq. (4)) for a constant initial speed *v*_t,start_. We left the four parameters (*K, τ, v*_t,start_, *v*_t,target_) free to vary in the fit. For the onset speed traces corresponding to continuous swimming, the best fit estimates of *v*_t,target_ matched the values to which *v*_*t*_ converged after the initial transient. The range of *v*_t,target_ obtained in this continuous case was used to constrain the fitting procedure for the remaining onset speed signals corresponding to intermittent swimming.

Using the inferred parameters, we computed the gain-delay product *Kτ* and the speed ratio *v*_t,start_*/v*_t,target_ to locate each fish within the dynamical phase space (Fig. 4**b**).

### 4.8 Manipulation of sensory feedback

#### 4.8.1 Vestibular: high-viscosity medium

Water viscosity was increased using dextran (Sigma-Aldrich, molecular weight 450 kDa to 650 kDa), prepared as an 8% (w/v) solution in water [49]. Viscosity was measured with a rheometer (Anton Paar), showing a tenfold increase compared to water at the same experimental temperature. The preparation and measurement were repeated three times, yielding consistent results. We chose to increase medium viscosity tenfold to match the nearly tenfold rise in Reynolds number observed from 2- to 5-week-old fish. This manipulation allowed us to artificially reverse the transition by mimicking the hydrodynamic changes experienced during development. These experiments were conducted in a small arena (7 *×* 7 cm^2^). The fish swimming was first monitored in water, then in the viscous medium (where it was kept for a maximum of 15 minutes), and finally back in normal water as a further control.

#### 4.8.2 Lateral line: ablation

The lateral line of four-week-old *Danionella* was chemically ablated with neomycin, following a protocol established in zebrafish larvae [50] and previously validated in adult *Danionella* [17]. Specifically, fish were immersed in a 200 µM neomycin (SigmaAldrich) solution for 30 minutes, then thoroughly rinsed in Ringer solution. Ringer solution was used to match blood osmotic pressure, which helped preserve cell integrity and function at the ablation site. The fish was then allowed to recover for 1-hour in the incubator. All experiments were carried out within 5 hours of the lateral line ablation to prevent neuromast regeneration.

Following Veith *et al*. [17], ablation was confirmed by exposing treated *Danionella* to a 50 µM solution of 4-Di-2-ASP (Sigma-Aldrich) for 5 minutes, followed by fluorescence imaging using a binocular microscope (see Fig. S10d). The 4-Di-2-ASP bulk solution was prepared in E3 medium with 1% ethanol to aid dissolution.

#### 4.8.3 Visual: complete darkness experiments

Experiments were conducted in complete darkness to deprive the fish of any visual cues: the projector was turned off, and the enclosure surrounding the entire setup was internally lined with black sheets and fully closed.

### 4.9 Virtual reality experiments

The experiments were conducted as described in Demarchi *et al*. [15]. The fish were presented with visual currents in the forward direction with an optic flow rate *ω* = 0.5 rad/s, lasting for 30 s, and separated by pauses of 30 s. We probed 5 different values of the feedback gain (the scaling factor *α* between fictive fish speed *v*_f_ and feedback optic flow rate *ω*), corresponding to 5 different target speeds *v*_f,target_ = *ω/α* for which the fish stabilizes its position in the virtual environment. Each target speed was presented in two distinct trials in a randomized fashion. To reduce the potential bias due to struggling movements, that are frequent in head-tethered configurations, the average bout duration per trial was computed as the expected duration of a swimming event selected by picking at random a time where the fish was swimming.

## 5 Supplementary information

## Abbreviations

BMI: Body mass index
COPD: Chronic obstructive pulmonary disease
CV: Coefficient of variation
DET: Percentage of determinism
EMD: Empirical mode decomposition
FEV_1_: Forced expiratory volume in the first second
LMM: Linear mixed models
MD: Mean difference
MMSE: Mini-mental state examination
MVC: Maximal voluntary contraction
MVF: Maximal voluntary force
PwCOPD: People with chronic obstructive pulmonary disease
RMSE: Root-mean-square error
ROF: Rating-of-fatigue scale
RP: Reccurence plot
RQA: Recurrence quantification analysis
SampEn: Sample entropy
SD: Standard deviation

## Acknowledgements

We thank the members of the IBPS aquatic animal facility for their assistance in animal care, and Doga Demirdelen for help with egg collection and larval rearing. We are grateful to Benjamin Judkewitz and Filippo Del Bene for sharing their *Danionella* lines, and to Julie Lafaye for helping setting up the *Danionella* facility. We also thank Ruben Portugues for valuable discussions, and Mattéo Dommanget-Kott for support with debugging.

## Funding

The European Union’s Horizon 2020 research and innovation programme under the Marie Sklodowska-Curie grant agreement No. 860949; ANR (Locomat project, ANR-21-CE16-0037).

## Author contribution

Conceptualization: M.C., G.D., L.D.; Data curation: M.C.; Formal analysis: M.C., L.D.; Funding acquisition: V.B., G.D.; Investigation: M.C., L.D., R.W.; Methodology: M.C., L.D., R.W., V.D., C.C., T.P., G.M.-D. G.G., V.B., G.D.; Project administration: G.D.; Resources: G.M.-D., V.B., G.D.; Software: M.C, V.D., R.W., L.D., T.P.; Supervision: G.D.; Validation: M.C.; Visualization: M.C., L.D., R.W.; Writing – original draft: M.C.; Writing review and editing: M.C., G.D., L.D., V.B., R.W., G.G.

## Competing interests

The authors declare no competing interests.

## Data and materials availability

Data and code are available on Zenodo.

## Notes

### Competing Interest Statement

The authors have declared no competing interest.

