## Supplementary Materials for "A sensorimotor instability drives a locomotor transition during fish development"

### 1 Supplementary text

#### 1.1 Statistical analysis of the ON and OFF states

To characterize how locomotor dynamics evolve, we analyzed the distributions of ON and OFF state durations (Fig. S2**b**).

Focusing first on the ON state durations, we found that young fish exhibit a broad distribution with a long tail extending up to several seconds, indicating prolonged swimming periods. As fish mature, this distribution becomes progressively narrower and develops a distinct peak, revealing the emergence of a characteristic bout duration whose mean value progressively shortens to settle at 240 ± 60ms (see Tab.S1 for summary of the statistics).

At early developmental stages, both $\tau_{\mathrm{ON}}$ and $\tau_{\mathrm{OFF}}$ follow an approximately exponential distribution. This indicates that, once the animal starts/stops swimming, the probability of stopping/starting again is constant over time, a hallmark of Poisson-like (memoryless) behaviors.

Regarding the OFF periods (Fig. S2**c**), at later stages, the distribution becomes peaked, with a characteristic idling time that increases with age to reach 430 ± 110ms. One should notice that at all ages, the distribution vanishes at finite time (of order 1s), which shows that the animals are engaged in swimming for the entire duration of the experiment.

| mean ± std | 1-2 wpf | 3 wpf | 4-5 wpf |
|---|---|---|---|
| $\bar{\tau}_{\mathrm{ON}}$ (ms) | 1190 ± 655 | 450 ± 320 | 240 ± 60 |
| $\bar{\tau}_{\mathrm{OFF}}$ (ms) | 370 ± 120 | 250 ± 65 | 430 ± 110 |

**Table S1** Mean ± standard deviation across fish (N = 4, 11, 5) of the individual median values of $\tau_{\mathrm{ON}}$ and $\tau_{\mathrm{OFF}}$.

#### 1.2 Analysis of linear delay equation

Here we analyze the behavior of Eq. (4). We denoted the target speed as $V_{\mathrm{target}}$, its value depends on which variable $V$ refers to. For $V = v_{\mathrm{f}}$, the target is simply $V_{\mathrm{target}} = v_{\mathrm{f,target}}$. For $V = v_{\mathrm{t}}$, it is $v_{\mathrm{t,target}} = v_{\mathrm{f,target}}/k_{\mathrm{f}}$. For $V = u$, it is $u_{\mathrm{target}} = v_{\mathrm{f,target}}/(k_{\mathrm{f}}k_u)$.

We have seen that the behavior of Eq.(4) depends on the value of the dimensionless product $K\tau$. To understand the behavior of the equation following th onset of swimming, we can consider a fish that starts swimming at speed $V_{\mathrm{start}}$ and, after a time $\tau$, regulates its speed based on the discrepancy between $V_{\mathrm{start}}$ and the target $V_{\mathrm{target}}$. If $K\tau < 1/e$ the speed converges to the target and the fish keeps swimming continuously (Fig. 4c first row). If $K\tau > \pi/2$ the speed diverges from the target, once it reaches zero the swimming event terminates, resulting in a finite bout (Fig. 4c third row). In the intermediate regime $1/e < K\tau < \pi/2$ the behavior will depend on the value of the initial speed. If $V_{\mathrm{start}}$ is close enough to the target the subsequent speed oscillations do not reach zero and the fish swims continuously (Fig. 4c second row). If $V_{\mathrm{start}}$ is large enough then the first undershoot of the target can reach zero, terminating the swimming event (Fig. 4c fourth row).

Eq. (4) with the initial condition $V(t) = V_{\text{start}}$ for $t \in [0, \tau]$ can be solved by integrating it iteratively with the method of steps [51]. For $V_{\text{start}} = 1$ and $V_{\text{target}} = 0$ one finds the following solution:

$$x(t) = \sum_{n=0}^{\infty} \left[ \frac{1}{n!} [-K(t - n\tau)]^n H(t - n\tau) \right] \tag{S1}$$

Where $H$ denotes the Heaviside step function. Therefore the solution for different values of $V_{\text{start}}$ and $V_{\text{target}}$ becomes:

$$V(t) = V_{\text{target}} + (V_{\text{start}} - V_{\text{target}})x(t) \tag{S2}$$

Eq. (S2) is what is shown in Fig. 4c for different values of the parameters.

Now, to determine whether the fish will swim continuously or terminate swimming in a finite time, we have to look at whether $V(t)$ reaches zero or remains positive for all times. We can do that by looking at the sign of its minimum $V_{\text{min}} = \min_t V(t)$:

$$\frac{V_{\text{min}}}{V_{\text{target}}} = 1 + \min_t \left( \frac{V_{\text{start}}}{V_{\text{target}}} - 1 \right) x(t) \tag{S3}$$

As varying $\tau$ corresponds to a rescaling of time, the minimum and maximum of $x(t)$ will only depend on the product $K\tau$. The value of $V_{\text{min}}$ is shown in Fig. 4b as a function of $K\tau$ and $V_{\text{start}}/V_{\text{target}}$. As we had anticipated, we find a region of the parameter space corresponding to continuous swimming (blue region in Fig. 4b) and one corresponding to discrete swimming (red region in fig. 4b).

For $K\tau > \pi/2$ one does not find a well-defined minimum because $V(t)$ can reach arbitrarily small values as $t \to \infty$. For $K\tau < 1/e$ one finds $V_{\text{min}} = V_{\text{target}}$ for $V_{\text{start}} > V_{\text{target}}$ and $V_{\text{min}} = V_{\text{start}}$ for $V_{\text{start}} < V_{\text{target}}$. Finally for $1/e < K\tau < \pi/2$ we find a transition from positive to negative values of $V_{\text{min}}$ as one increases the starting speed. The boundary between the two regions can be found by setting $V_{\text{min}} = 0$ in Eq. (S3):

$$\left. \frac{V_{\text{start}}}{V_{\text{target}}} \right|_{V_{\text{min}}=0} = 1 - \frac{1}{x_{\text{min}}(K\tau)} \tag{S4}$$

Thus, one sees that as $K\tau$ increases $x_{\text{min}}$ decreases and one can reach a negative $V_{\text{min}}$ for progressively smaller values of $V_{\text{start}}/V_{\text{target}}$. Eq. (S4) holds for $V_{\text{start}} > V_{\text{target}}$, the analogous curve for $V_{\text{start}} < v_{\text{target}}$ is obtained by considering the $x_{\text{max}}$ instead of $x_{\text{min}}$. In the case of $V_{\text{start}} > V_{\text{target}}$, one finds that $V_{\text{min}}$ is almost everywhere equal to $V_{\text{start}}$, except for a small region for $K\tau$ close to $\pi/2$ where the first minimum of $V(t)$ falls below $V_{\text{start}}$, and, for small values of $V_{\text{start}}/V_{\text{target}}$, it can fall below zero. Overall, the boundary defined by $V_{\text{min}} = 0$ is plotted as a black dashed line in Fig. 4b.

#### 1.3 Analysis of nonlinear delay equation

Here we consider the nonlinearities resulting from logarithmic coding and look at how they change the model and the analysis. In this case, the motor command encodes the tail speed logarithmically $v_\mathrm{t} = A_\mathrm{t} e^{k_\mathrm{t} u}$ and it is updated based on a logarithmic representation of the fish speed $\dot{u} = -k_u[\log(v_\mathrm{f}) - \log(v_\mathrm{f,target})]$ (the corresponding time delays are implied). Then, the equation for the speed, both for $v_\mathrm{t}$ and $v_\mathrm{f}$, becomes:

$$\frac{dv}{dt} = -k_\mathrm{t} k_\mathrm{u} v(t) \log\left(\frac{v(t-\tau)}{v_\mathrm{target}}\right) \tag{S5}$$

Where $v_\mathrm{target} = v_\mathrm{f,target}$ for $v = v_\mathrm{f}$ and $v_\mathrm{target} = v_\mathrm{t,target}$ for $v = v_\mathrm{t}$. The equation for the motor command remains analogous to Eq. (4):

$$\frac{du}{dt} = -k_\mathrm{t} k_\mathrm{u}[u(t-\tau) - u_\mathrm{target}] \tag{S6}$$

Thus we expect the same analysis to hold for the dynamics of the motor command, but the speed $v$ is not simply proportional to $u$, but it it obtained through an exponential transformation. Then, $v$ is bound to have positive values, but it can get arbitrarily close to zero, as $\log(v)$ diverges to $-\infty$. We can expect the fish to stop swimming when $v$ falls below a certain threshold value below which it cannot swim. The exact behavior of the model depends on the choice of this threshold, but it remains qualitatively similar to that of the linear equation, where we simply chose the threshold to be zero.

For visuomotor stabilization against external currents it was shown that the presence of logarithmic nonlinearities moves the dependence of the system stability from the feedback gain to the intensity of the optic flow [15]. We note that here the situation is different, as the stability behavior of Eq. (S5) is the same as that of Eq. (S6), with the dimensionless parameter combination being $k_\mathrm{t} k_\mathrm{u} \tau$. This can be easily seen by noting that one equation is mapped to the other under the transformation $u = \log(v)$. What changes is that the target is internal (difference between logarithmic representation of perceived and target speed), whereas for the stabilization of an external current the target is external (difference between external and feedback optic flow), leading to a different functional form of the underlying equation.

This difference explains why we can observe old fish swimming continuously in head-fixed conditions, even though our model for spontaneous swimming (Eq. (4)) predicts that they would always swim intermittently, as they fall in the unstable regime. If we consider the model resulting for an external target in the presence of nonlinearities, we find that it is always stable, with the solution either converging to a fixed point or to a limit cycle [15]. Then, one can observe both intermittent and continuous swimming, depending on whether the swimming speed falls below a minimum threshold or not.

### 2 Supplementary figures

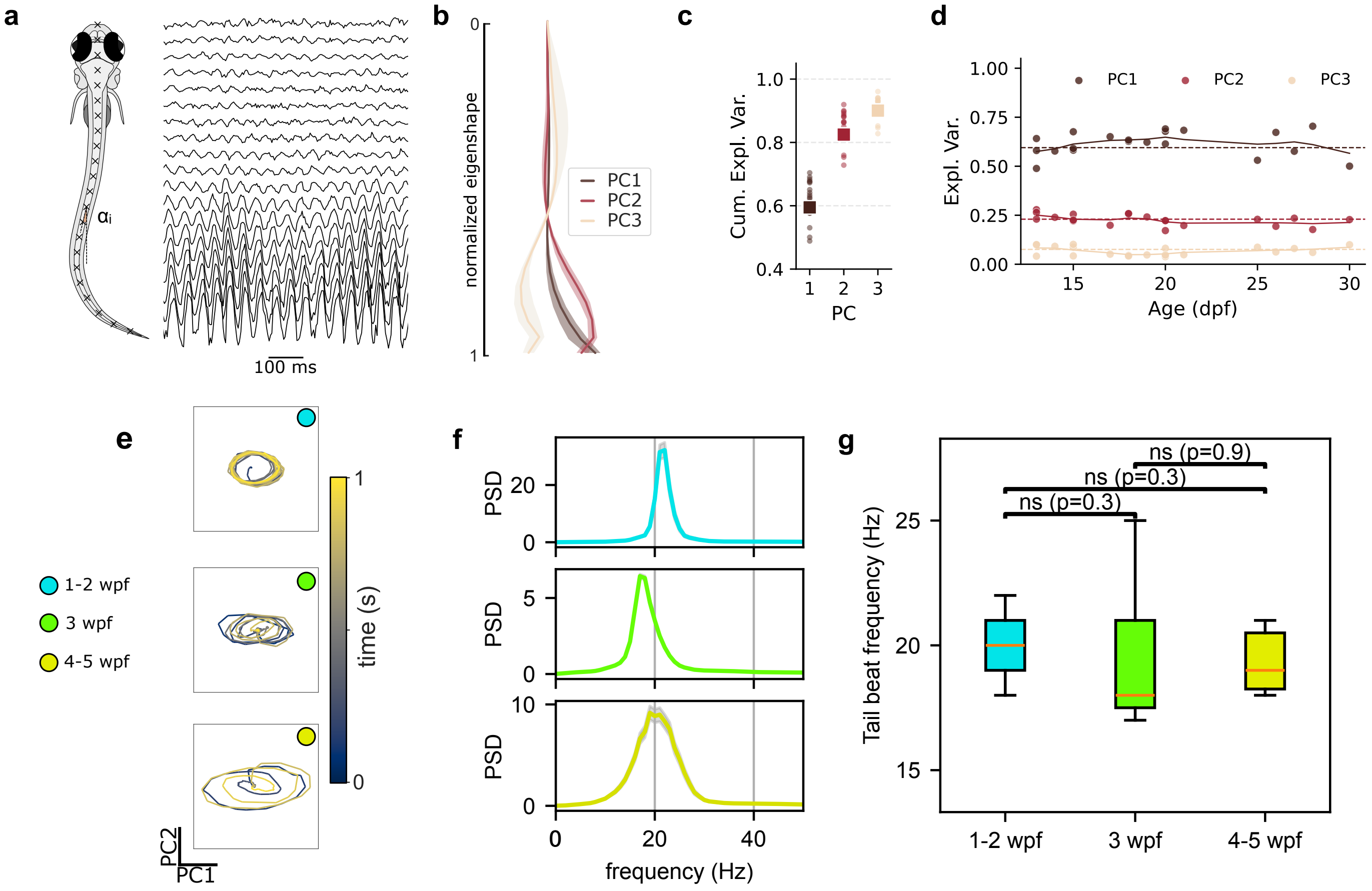


**Fig. S1 Conserved tail kinematics across development in *Danionella*. a**, Fish midline segmented into 20 points from head to tail; posture described by angles $\alpha_i$ relative to the body axis. PCA was applied to length-normalized $\alpha_i$. **b**, First three eigenshapes (bold); shaded areas show s.d. across fish. **c**, Cumulative explained variance of first three PCs. Dots: individual fish; squares: pooled data. **d**, Explained variance of first three PCs vs. age (dpf). Dots: individuals; solid line: smoothed trend; dashed line: pooled PCA values (59.49–22.98–7.57%). Variance remains stable across development. **e**, Dynamics in shared postural space (length-normalized PCA). **f**, PSD of tail deflection for fish at 13, 22, 28 dpf. Solid lines: mean PSD; shaded areas: s.e.m. **g**, Tail beat frequency distributions across ages. No significant differences (t-test, $p > 0.05$).

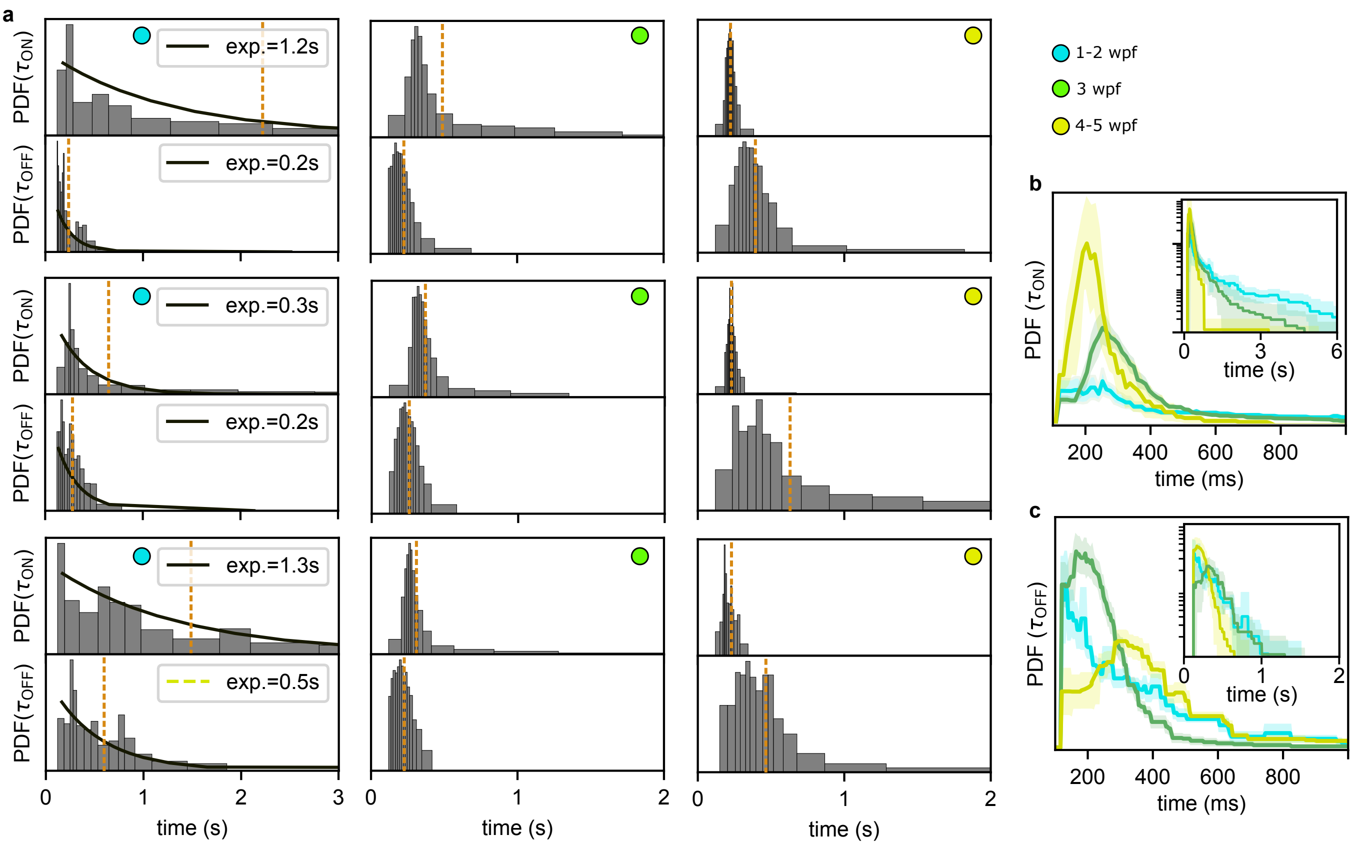


**Fig. S2 Statistics of $\tau_{ON}$ and $\tau_{OFF}$. a**, Constant-count histogram of $\tau_{ON}$ and $\tau_{OFF}$ for representative individual fish randomly selected from each of the three groups. For the young group, we fitted an exponential decay to highlight the absence of a characteristic timescale that emerge later in development and is a signature of intermittent swimming; both $\tau_{ON}$ and $\tau_{OFF}$ follow a Poisson-like distribution at this stage. **b-c**, Distribution of $\tau_{ON}$ and $\tau_{OFF}$. The solid lines and shaded areas represent the mean and standard deviation across fish grouped by age (blue: 1–2 wpf (N=4), green: 3 wpf (N=11), yellow: 4–5 wpf (N=5)). Inset: the same distributions shown on a semi-logarithmic (log–y) scale to facilitate visualisation of the long-time tail.

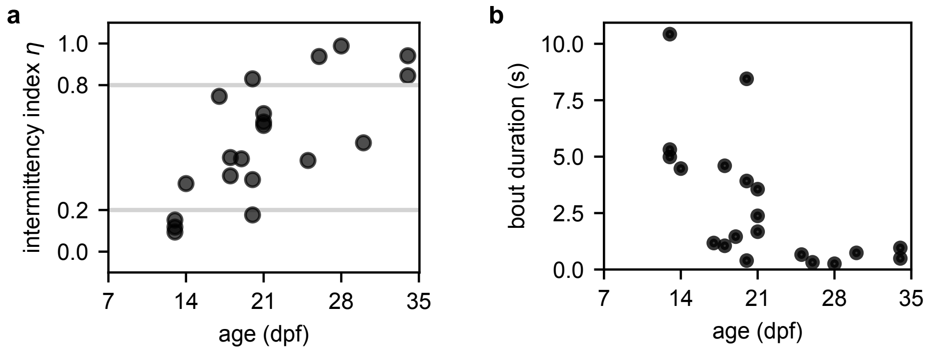


**Fig. S3 Developmental changes in intermittency index and bout duration a**, Intermittency index $\eta$ plotted for each fish as a function of age. The corresponding boxplot grouped by age is shown in Fig. 1.**h** in the main text. **b**, Bout duration for each fish as a function of age. It is computed as the expected duration of the swimming period for a given time during which the fish is swimming.

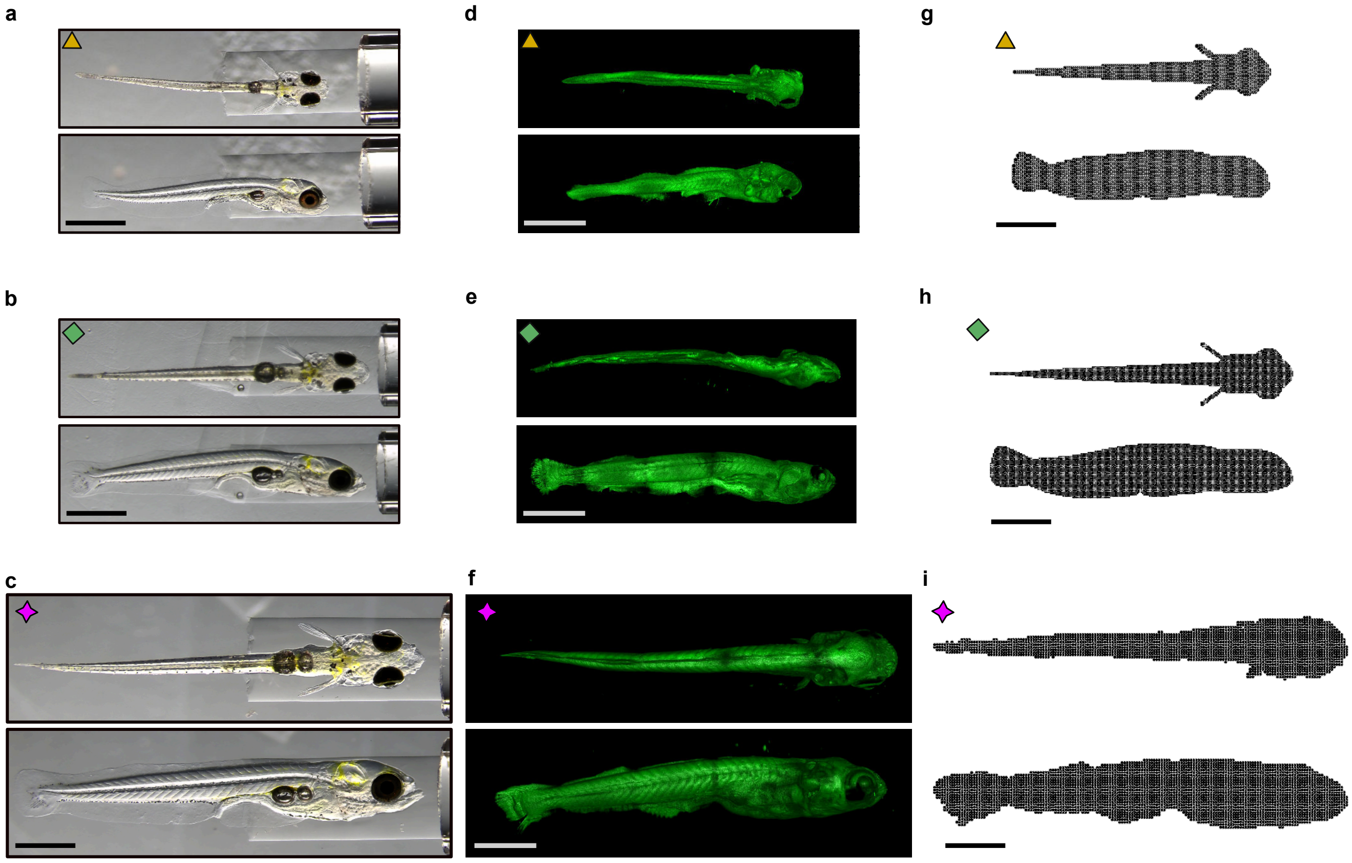


**Fig. S4 Building 3D models of Danionella used for CFD simulations across ages. a-b-c**, Dorsal (top) and frontal (bottom) views of 2D images of head-tethered *Danionella* at 14 dpf (a), 23 dpf (b), and 30 dpf (c). For (a) and (c), the same images are presented in the main Fig. 5 and are reproduced here for completeness. Scale bar: 1 mm. **d-e-f**, Dorsal (top) and frontal (bottom) views of raw 2D lightsheet scans stitched to reconstruct the 3D morphology of *Danionella*. See Methods for details on the clearing, lightsheet scanning and stitching procedure. Scale bar: 1mm. **g-h-i** Final 3D models used for CFD simulations, shown as point clouds. The intermediate-age fish (h) is used solely for calibration. Scale bar: 1 mm.

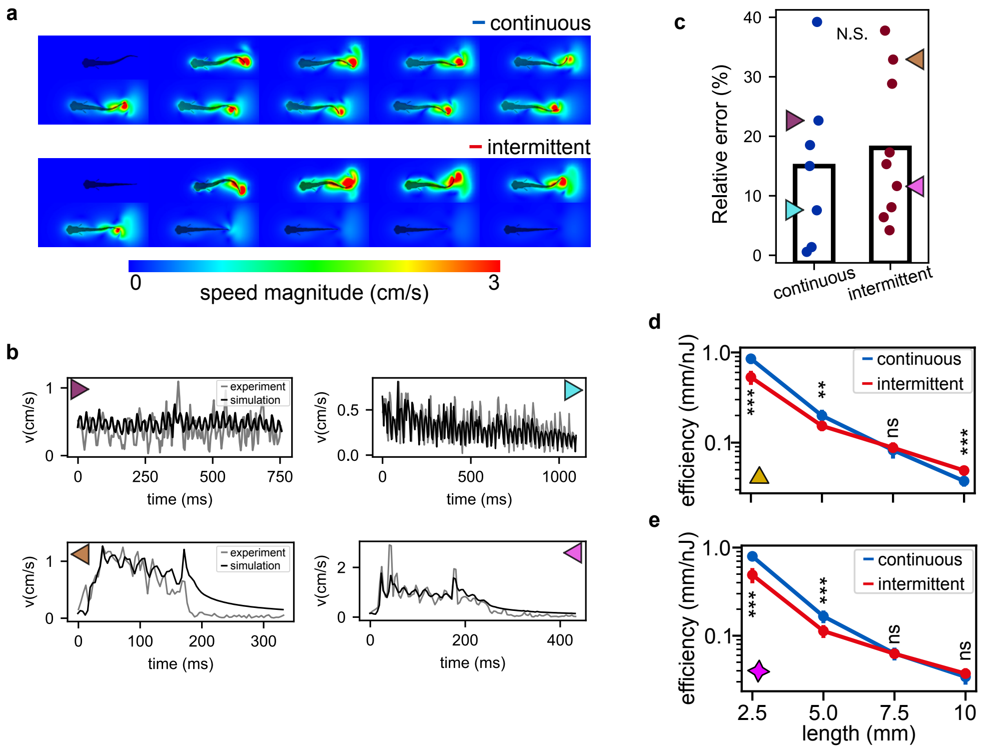


**Fig. S5 Calibration of continuous and intermittent swimming modes using intermediate-age fish.** **a**, Representative snapshots of continuous (top) and intermittent (bottom) swimming, spaced 38 ms apart. **b**, Example comparisons between estimated and simulated speeds for the two swimming modes. Top: continuous swimming, excluding an initial 0.5-second “burn-in” period to allow simulated speed to match experimental initial conditions. Bottom: burst-and-glide swimming. Colored arrows indicate specific data points shown in panel (c). **c**, CFD simulations show minimal relative error compared to the average experimental speeds, with no statistically significant difference (two-sample t-test). Each dot represents a 0.7–2.5s continuous swimming segment or a single burst-and-glide bout. Colored arrows correspond to the comparisons shown in panel (b). **d**, Efficiency vs. body length using the 14 dpf *Danionella* morphology. Mean ± std across simulated events for continuous (blue, N = 11, 11, 11, 9) and intermittent (red, N = 14, 14, 14, 11) swimming. A crossover in energetic efficiency occurs near $L \sim 7$ mm. At $L = 7.5$ mm, the difference is not significant ($p = 0.4$); at other lengths, it is: $L = 2.5$ mm ($p = 10^{-7}$), 5 mm ($p = 10^{-3}$), and 10 mm ($p = 10^{-4}$) (t-test). **e**, Same analysis as in (f) using the 30 dpf morphology. Continuous (blue, N = 11, 11, 11, 9) and intermittent (red, N = 14, 14, 12, 13) swimming show a crossover near $L = 7.5$ mm ($p = 0.6$); significant differences occur at 2.5 mm ($p = 10^{-8}$) and 5 mm ($p = 10^{-5}$), while at $L = 10$ mm the difference is not significant ($p = 0.3$) (t-test).

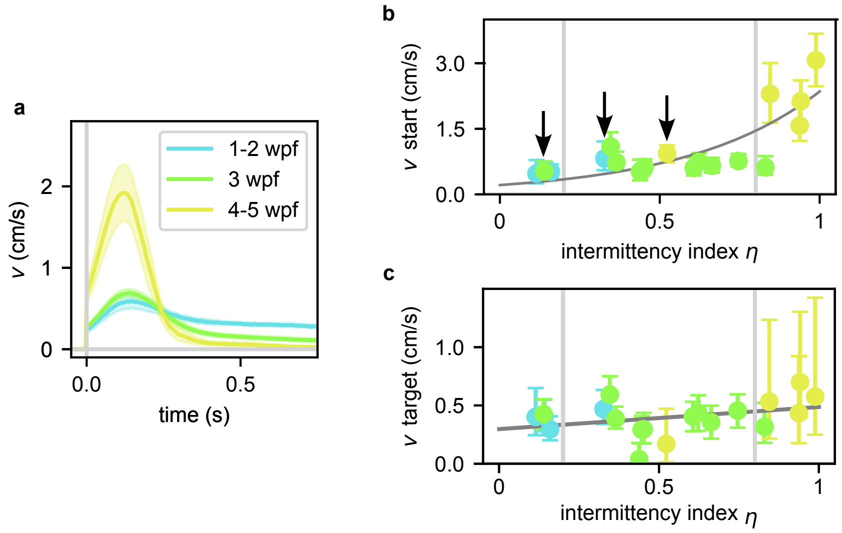


**Fig. S6 Onset speed as functions of age and intermittency index $\eta$. a**, Fish speed aligned to movement onset for straight swimming events. Here, fish were grouped according to their age, in contrast to Fig. 3**a** of the main text where we used the intermittency index $\eta$. Bold lines represent the mean across fish; shaded regions indicate the standard deviation. **b**, Maximum onset speed plotted against the intermittency index $\eta$. Each point represents the median and interquartile range for an individual fish, colour-coded by age group. Arrows indicate fish whose $\eta$ value correlates more strongly with $v_{start}$ than with chronological age. The grey curve is the exponential best-fit to the data, shown as a visual guide. **c** Target speed plotted against the intermittency index $\eta$. $v_{target}$ is defined as the temporal average of each swimming event. The grey curve represents a linear best-fit to the data, shown as a visual guide.

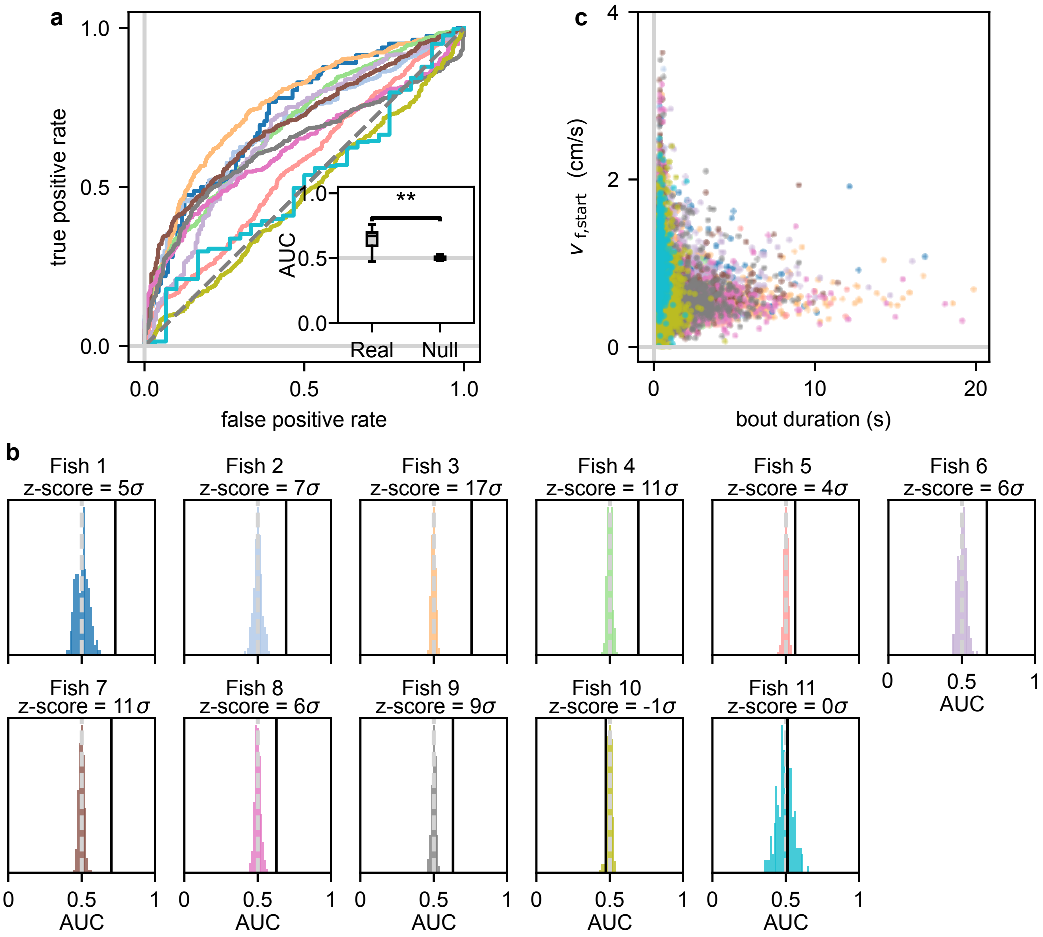


**Fig. S7 Testing the predictive power of $v_{f,start}$ on the bout outcome. a**, Receiver Operating Characteristic (ROC) analysis assessing whether $v_{f,\mathrm{start}}$ predicts the outcome of swimming events, binarized as either continuous or intermittent. The 'Real' ROC curves are shown for each fish in different colours. The area under the curve (AUC) is compared with the AUC from 'Null' ROC curves, generated by randomizing both $v_{f,\mathrm{start}}$ and the labels (inset). A non-parametric Mann–Whitney U test reveals a significant difference (p = 0.009). **b**, To quantify fish-specific variability, we ran the 'Null' model 1000 times and show the resulting distribution. The z-score between its mean and the real AUC is also reported. **c**, $v_{f,\mathrm{start}}$ as a function of bout duration. Lower speeds tend to result in longer swimming bouts, while higher speeds are associated with shorter events. A dense cluster of points at low speeds and short durations may reflect mislabeling errors, as thresholding becomes less reliable at low speeds. Data from all fish are shown, with each individual represented by a different colour. Due to the high throughput of the experiment, a large number of swimming events were recorded per fish: on the order of $10^2$ for continuous events and $10^3$ for bout-like events.

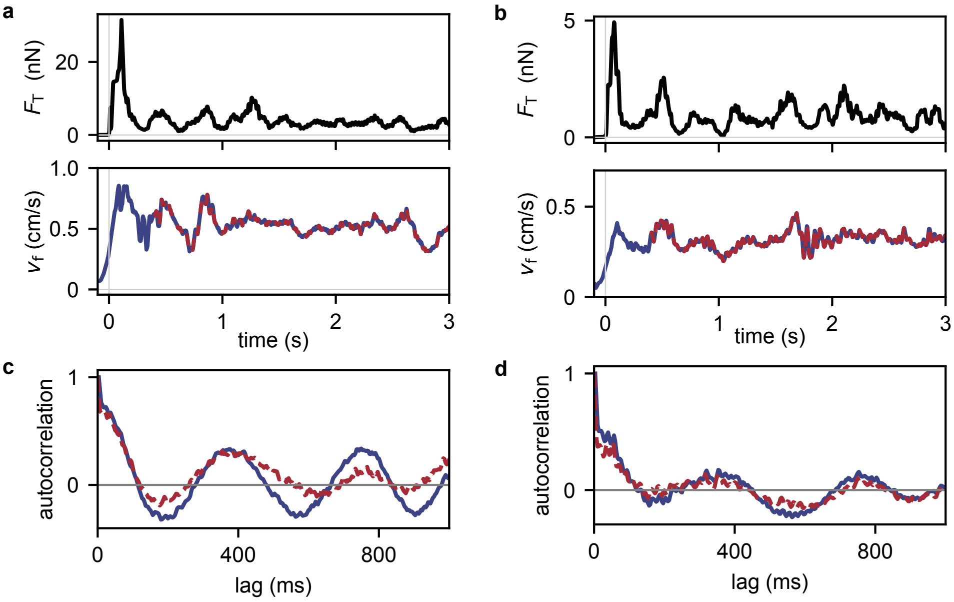


**Fig. S8 Example of speed modularion during long swimming events. a-b**, Examples of thrust signals (top, $F_T$) and fish speed (bottom, $v_f$) during two prolonged swimming events (first 3 seconds shown) in two different fish of 17 dpf (**a**) and 21 dpf (**b**). Sustained speed oscillations were observed throughout the duration of the swimming event. **c-d** Characterization of speed oscillations using autocorrelation analysis. The autocorrelation of the full signal (blue) is compared with that of the signal excluding the first 0.4 s (red). The latter isolates the characteristic modulation frequency from the initial overshoot. The two autocorrelation curves overlap closely. The typical period of those speed oscillations is approximately 400ms.

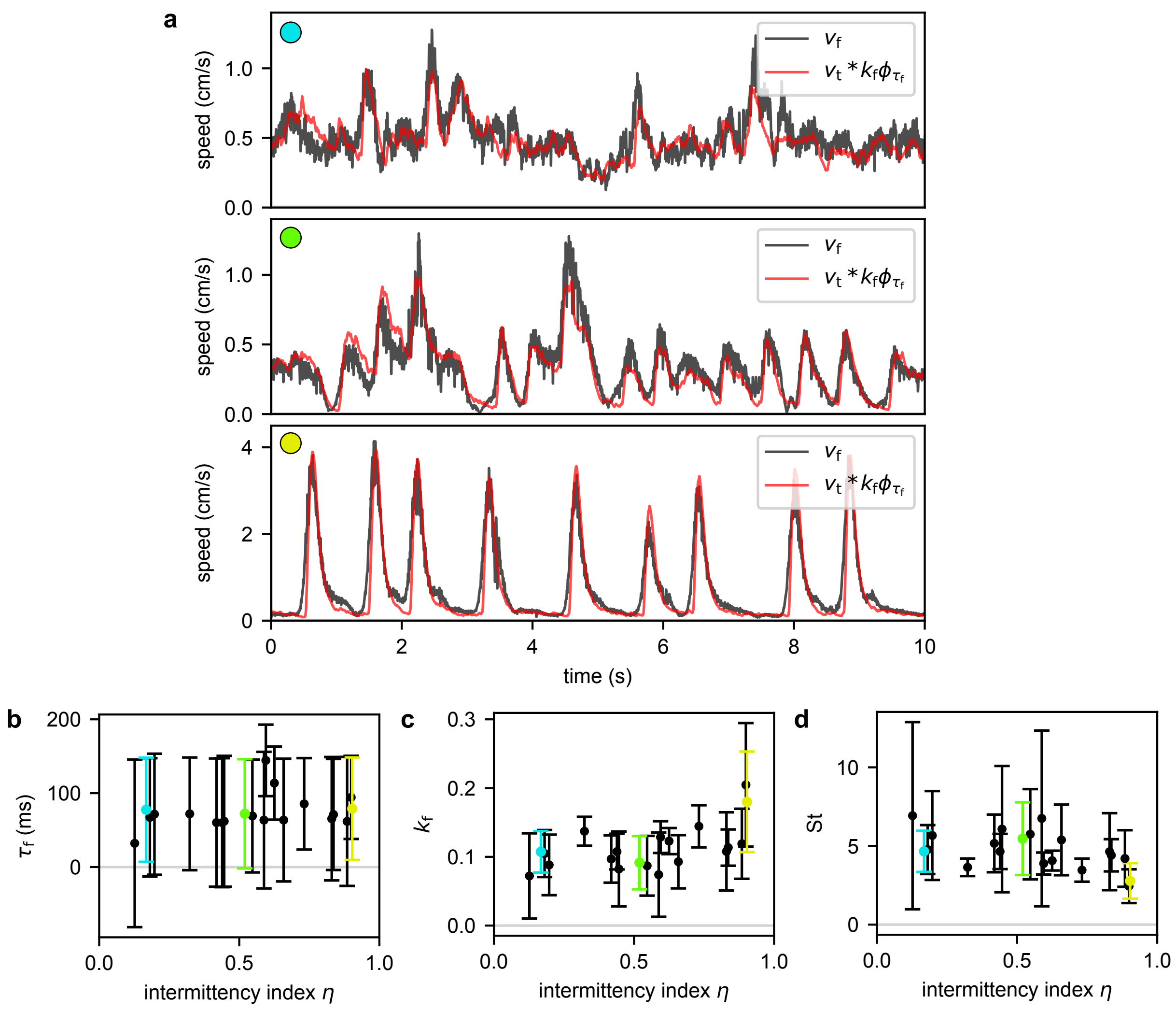


**Fig. S9 Relationship between tail speed and fish speed.** **a**, Traces comparing the fish (centre-of-mass) speed $v_\mathrm{f}$ with the delayed tail speed: $k_\mathrm{f} v_\mathrm{t} * \phi_{\tau_\mathrm{f}}$, where $\phi_{\tau_\mathrm{f}} = e^{-t/\tau_\mathrm{f}}/\tau_\mathrm{f}$ is an exponential temporal kernel, and $k_\mathrm{f}$ is a rescaling factor. The comparison of the two signals is shown for three representative fish corresponding to the three groups (top: 1–2 wpf, center 3 wpf, bottom 4–5 wpf). The values of $k_\mathrm{f}$ and $\tau_\mathrm{f}$ were infered by minimizing the mean squared error between the two signals. **b**, Estimated delay $\tau_\mathrm{f}$ between tail and fish speed, as a function of the intermittency index $\eta$. Each data point represents the mean and standard deviation for one fish, weighted with the duration of each uninterrupted recording sequence. The values for the example fish shown in (**a**) are colored with the corresponding subplot labels. **c** Analogous to (**b**), but for the scaling factor $k_\mathrm{f}$ between tail and fish speed. **d** Analogous to (**b**), but for the Strouhal number $\mathrm{St} = 1/(2k_\mathrm{f})$.

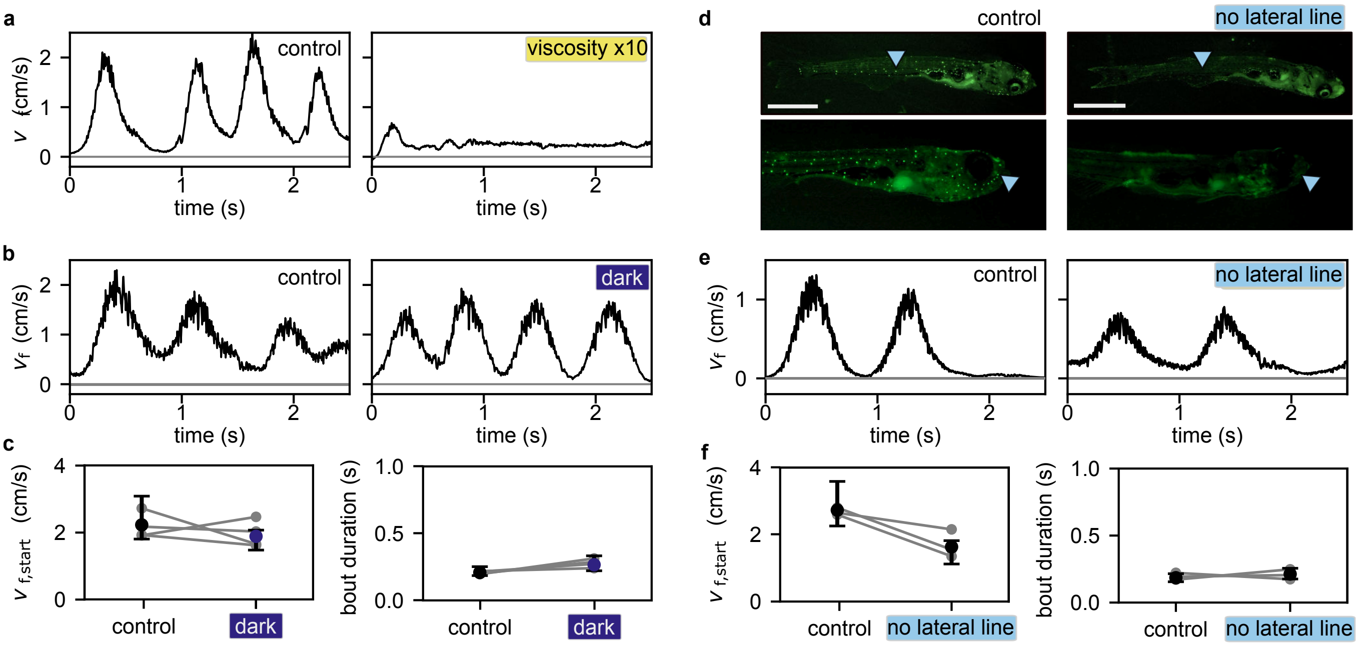


**Fig. S10 Effects of externally manipulating vestibular, visual, and lateral line inputs in freely swimming *Danionella*.** **a**, Example traces of swimming speed in 5 wpf *Danionella* under control conditions in water (left) and in a viscous medium (right). **b**, Example traces of swimming speed in 5 wpf *Danionella* under control light conditions (left) and in complete darkness (right). **c**, $v_{\text{start}}$ and bout duration (median ± IQR) across fish (N = 5) in dark and control conditions; grey lines indicate mean values of swimming events for individual fish. **d**,*Danionella* stained with DASPEI, imaged with an epifluorescence microscope to control lateral line ablation. Scale bar: 1 mm. **e**, Example traces of swimming speed in 4 wpf *Danionella* under control conditions (left) and after lateral line ablation (right). **f**, $v_{\text{start}}$ and bout duration (median ± IQR) across fish (N = 3) in lateral line ablated fish and control conditions; grey lines indicate mean values of swimming events for individual fish.

### 3 Supplementary movies

**Movie S1: Freely swimming 14 dpf Danionella in the experimental arena.** Multiple timescales are illustrated: at the millisecond scale, tail kinematics are segmented (movie slowed 10× to 25 FPS); at the second scale, the fish swims continuously, exhibiting ON/OFF dynamics; and at the minute scale, it explores the entire arena.

**Movie S2: Freely swimming 33 dpf Danionella in the experimental arena.** Multiple timescales are illustrated: at the millisecond scale, tail kinematics are segmented (movie slowed 10× to 25 FPS); at the second scale, the fish swims intermittently, exhibiting ON/OFF dynamics; and at the minute scale, it explores the entire arena.

**Movie S3: CFD simulations of intermediate-age fish swimming continuously.** *Left*: Continuous swimming event from the raw video of a 23 dpf freely swimming *Danionella*, slowed 10× to 25 FPS. The relative tracked fish spline was used to animate the intermediate-age 3D *Danionella* model. *Right*: Velocity field extracted from the CFD simulation. Scale bar: 1 mm.

**Movie S4: CFD simulations of intermediate-age fish swimming intermittently.** *Left*: Intermittent swimming event from the raw video of a 23 dpf freely swimming *Danionella*, slowed 10× to 25 FPS. The relative tracked fish spline was used to animate the intermediate-age 3D *Danionella* model. *Right*: Velocity field extracted from the CFD simulation. Scale bar: 1 mm.

**Movie S5: CFD simulations of two weeks old fish.** Velocity field for a continuous swimming event from a 13 dpf *Danionella*. Simulation performed using the 14 dpf morphology.

**Movie S6: CFD simulations of four weeks old fish.** Velocity field for a burst-and-glide bout from a 34 dpf *Danionella*. Simulation performed using the 30 dpf morphology.

**Movie S7: Four weeks old fish swimming in high viscosity medium.** *Left*: 32 dpf *Danionella* freely swimming in water. *Right*: the same fish swimming in a high-viscosity medium. Movies are slowed 10x to 25 FPS. Scale bar: 1 mm.
